# Loss of *Zmiz1* in mice leads to impaired cortical development and autistic-like behaviors

**DOI:** 10.1101/2024.08.18.608498

**Authors:** K C Rajan, Nehal R Patel, Abbigail Thurmon, Maryann Grace Lorino, Alina S. Tiemroth, Isabella Kulstad, Vivianne Morrison, Mauren Akumuo, Anoushka Shenoy, Stryder M Meadows, Maria J Galazo

## Abstract

De novo mutations in transcriptional regulators are emerging as key risk factors contributing to the etiology of neurodevelopmental disorders. Human genetic studies have recently identified ZMIZ1 and its de novo mutations as causal of a neurodevelopmental syndrome strongly associated with intellectual disability, autism, ADHD, microcephaly, and other developmental anomalies. However, the role of ZMIZ in brain development or how ZMIZ1 mutations cause neurological phenotypes is unknown. Here, we generated a forebrain-specific *Zmiz1* mutant mouse model that develops brain abnormalities, including cortical microcephaly, corpus callosum dysgenesis, and abnormal differentiation of upper-layer cortical neurons. Behaviorally, *Zmiz1* mutant mice show alterations in motor activity, anxiety, communication, and social interactions with strong sex differences, resembling phenotypes associated with autism. Molecularly, *Zmiz1* deficiency leads to transcriptomic changes disrupting neurogenesis, neuron differentiation programs, and synaptic signaling. We identified *Zmiz1*-mediated downstream regulation of key neurodevelopmental factors, including *Lhx2, Auts2*, and *EfnB2*. Importantly, reactivation of the EfnB2 pathway by exogenous EFNB2 recombinant protein rescues the dendritic outgrowth deficits in *Zmiz1* mutant cortical neurons. Overall, our in vivo findings provide insight into *Zmiz1* function in cortical development and reveal mechanistic underpinnings of ZMIZ1 syndrome, thereby providing valuable information relevant to future studies on this neurodevelopmental disorder.

## INTRODUCTION

Neurodevelopment is a tightly controlled multi-step process, including neuron generation, differentiation, wiring, and synaptic formation. Dysregulation at any point in this process may lead to neurodevelopmental disorders (NDDs). Within this complex landscape, transcription factors (TFs), transcription co-regulators, and chromatin remodelers have emerged as central elements orchestrating neurodevelopment, and as key risk factors for NDDs (1). However, the contribution of mutations, particularly de novo mutations, in these regulatory factors to NDDs remains poorly understood (2–6).

Recent human genetic studies have linked de novo, loss of function mutations in Zinc Finger MIZ-Type Containing 1 (ZMIZ1) to a neurodevelopmental syndrome strongly associated with intellectual disability (ID), autism (ASD), attention deficit and hyperactivity disorder (ADHD), microcephaly, and other developmental anomalies (6–11). Also, single nucleotide variations in a ZMIZ1 enhancer have been identified as a risk factor for ASD (6), further highlighting the link between ZMIZ, neurodevelopment, and ASD. *Zmiz1* functions as a transcriptional co-activator for important developmental pathways, including Notch1 (7), Androgen Receptor (8), p53 (9), and Smad3/4 (10), and interacts with proteins of the SWI/SNF-like BAF chromatin remodeling complex (12), which underscores its potential significance in brain formation. Despite ZMIZ1’s strong association with human NDDs, its role in neurodevelopment, and how ZMIZ1 mutations contribute to ASD risk and NDDs remains largely unknown.

The cerebral cortex plays a critical role in sensory processing, motor control, and higher-order cognitive functions including learning, memory, and cognition. Abnormal cortical formation and connectivity are associated with complex NDDs and neurological phenotypes (2,13). In humans and mice, *Zmiz1* expression is strong in the embryonic cortex and decreases postnatally (14). Its cortical expression, roles in transcriptional control, and strong link to NDDs suggest that *Zmiz1* plays an important role in cortical development; however, its function and molecular targets in the cortex have not been studied.

Here, we generated a forebrain-specific *Zmiz1* mutant mouse model to investigate the role of *Zmiz1* in cortical development, and the resulting cortical morphological alterations, behavioral consequences, and molecular pathways dysregulated upon its deletion. We find that loss of *Zmiz1* in the cortex results in cortical microcephaly through reduced cortical progenitor proliferation and ultimately fewer cortical neurons. Interestingly, *Zmiz1* loss leads to dysgenesis of the corpus callosum, which is a phenotype commonly observed in NDDs. Focusing on callosal projection neurons, we find that *Zmiz1* regulates differentiation of layer 2/3 neurons, and its deletion specifically affects apical dendritic morphology. Importantly, deletion of *Zmiz1* in mice results in behavioral alterations that recapitulate aspects of human neurodevelopmental phenotypes in ZMIZ1 human patients. The behavioral phenotypes are present from early postnatal stages to adulthood, have different manifestations in males and females, and include deficits in communication, motor activity, anxiety, sensory gating, and social interactions. We investigated the transcriptional landscape and molecular targets regulated by *Zmiz1* at multiple cortical developmental stages. Integrating transcriptomic analysis and *Zmiz1*-ChIPseq, we identify *EfnB2, Auts2, Lhx2, EphA5*, and *Unc5D* as direct transcriptional targets of *Zmiz1*. Moreover, we show that reactivation of the Eph-ephrin pathway via EFNB2-fc recombinant protein treatment rescues the neurite growth deficits observed in *Zmiz1* mutant cortical neurons.

Collectively, our study unveils crucial functions of *Zmiz1* in cortical development, and sheds light on the molecular mechanisms and pathways underlying ZMIZ1 developmental syndrome, thereby deepening our understanding of the complex interplay between molecular processes and neurodevelopmental disorders.

## RESULTS

### Loss of *Zmiz1* alters cortical progenitor proliferation and leads to cortical microcephaly

In adult mice, *Zmiz1* is expressed throughout the brain, though its expression is highest in the cortex and cerebellum. *Zmiz1* is expressed across cell types, including neurons, oligodendrocytes, microglia, astrocytes, and endothelial cells (14). Although previous studies identified *Zmiz1* as an early cortical developmental gene (15), its function in the cortex is completely unknown. To begin to understand its role in cortical development, we investigated *Zmiz1* expression using publicly available single-cell transcriptomic atlases of the developing cortex (16,17). *Zmiz1* mRNA is highly expressed during embryonic cortical development, specifically in apical and basal progenitors and postmitotic neurons (Supplementary Fig. 1A, B). *Zmiz1* expression is strong in the embryonic cortex and decreases postnatally (14, 15).

To investigate the developmental role of *Zmiz1* in the cortex, we utilized *Zmiz1* conditional loxP flanked mice (*Zmiz1*^*fl/fl*^) (98) and *Emx1-Cre* mice (99) to generate a *Zmiz1* loss-of-function mutation in forebrain-specific structures, including the cerebral cortex and hippocampus (*Emx1-Cre;Zmiz1*^*fl/fl*^ – referred to as *Zmiz1*-KO). Efficient deletion in the cortex was validated by ZMIZ1 immunostaining (Supplementary Fig. 2). Focusing on the cortex, we first assessed whether *Zmiz1* deletion led to gross morphological phenotypes by quantifying dorsal cortical surface area and cortical midline length at postnatal day (P) 1, P3, P7, P14, and P21. We found a significant decrease in cortical surface area at P7, P14, and P21 while midline length was significantly decreased at P1, P3, P7, P14, and P21 in *Zmiz1*-KO cortex compared to control mice (*Zmiz1*^*fl/fl*^) (Fig. 1A-C). To assess whether *Zmiz1* mutation also affects cortical thickness and layer formation during development, we performed immunostaining for TBR1, CTIP2, and CUX1, which are transcription factors key for the development of neurons in cortical layer 6 (L6), layer 5 (L5), and upper layers (L2/3 and L4), respectively. We assessed the overall cortical thickness, layer thickness, and the distribution and density of TBR1+, CTIP2+, and CUX1+ in each layer (Fig. 1D-H). While the layer pattern was not altered, there was a significant reduction in the overall cortical thickness, and the thickness of every layer in *Zmiz1*-KO cortices (Fig. 1E). When layer thickness was normalized to the overall cortical thickness, only L6 was significantly reduced (Fig. 1G), indicating that the *Zmiz1*-KO cortex is thinner overall compared to controls, but L6 is significantly more reduced. Furthermore, we quantified the number of CUX1+, CTIP2+, and TBR1+ neurons in regions of interest of a fixed width, spanning from pia to white matter (Fig. 1D). As expected, given the reduction in cortical thickness, the number of CUX1+, CTIP2+, and TBR1+ neurons per column were significantly decreased in *Zmiz1*-KOs (Fig. 1F), however, their density in upper layers, L5, and L6, respectively, was not significantly different (Fig. 1H). Overall, these results indicate that *Zmiz1* mutation results in smaller cortices with reduced surface area and thickness, and reduced number of neurons in all layers.

**Figure 1:**
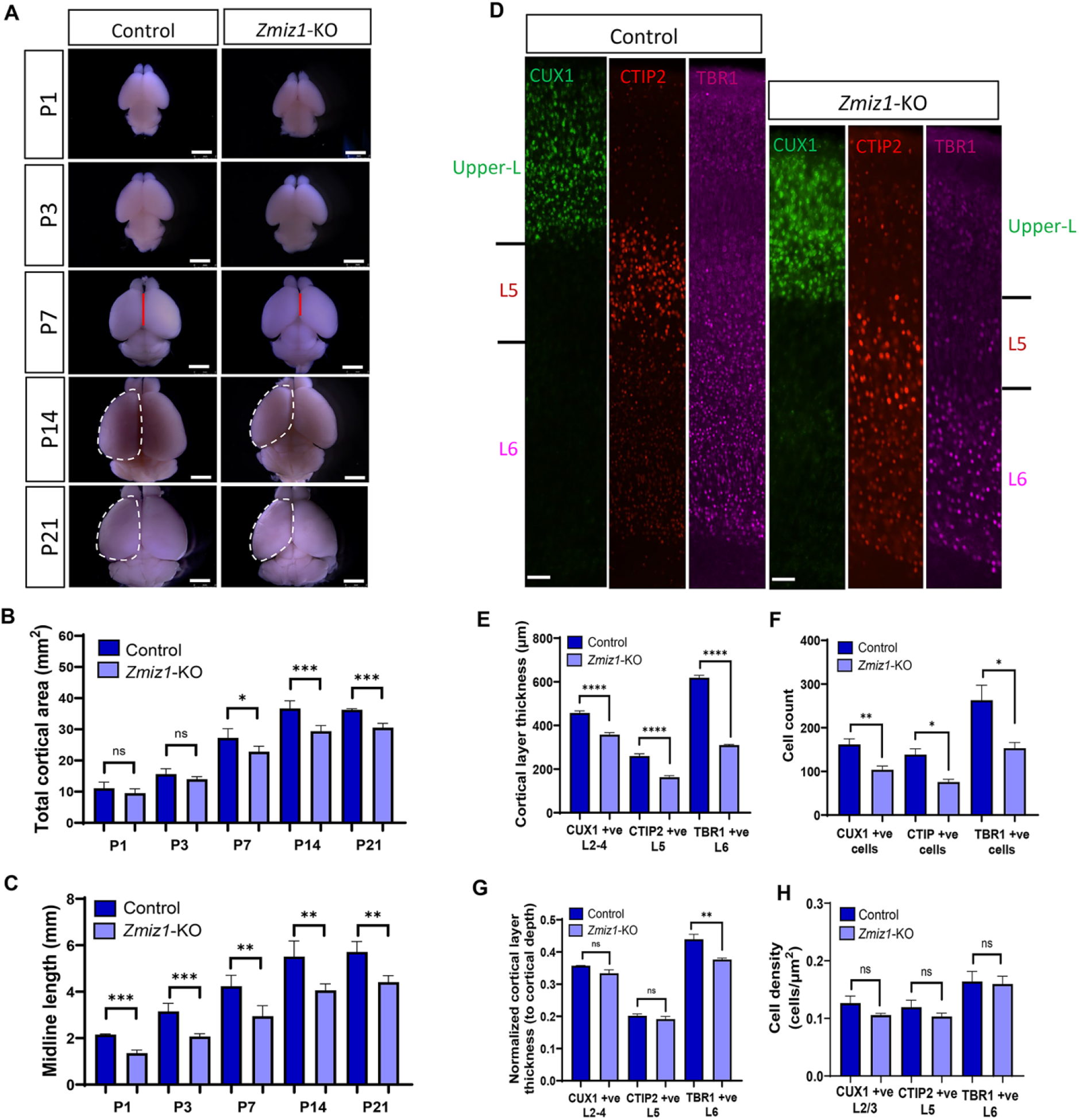
*Zmiz1* loss in the cortex results in cortical microcephaly. (A) Dorsal view of the brain showing reduced cortical area and midline length in *Zmiz1*-KO mice. (B-C) Quantification of (B) cortical area (example dashed line in P14 and P21), and (C) midline length (example, red line in P7) from P1 to P21. (D) Cortical layer immunolabeling using CUX1, CTIP2, and TBR1 at P7. (E) Quantification of cortical layer thickness. (F) Cell count of represented immunolabeled cells cortical layer. (G) Normalized cortical layer thickness to cortical depth. (H) Cell density for each cortical layer. Data are presented as mean ± SEM. * P <0.05, **P <0.01, ***P < 0.001, ****P < 0.0001, two-tailed unpaired t-test. n = 3-5 per group. Scale bars, 2 mm (A), 150 mm (D).

Malformations of cortical development such as microcephaly or megacephaly are associated with deficits in cortical neurogenesis (18). Given *Zmiz1* expression in progenitors and the cortical microcephaly phenotype observed in *Zmiz1*-KO mice, we hypothesized that *Zmiz1* may regulate cortical progenitor proliferation and neurogenesis. To investigate whether cortical progenitors are affected in *Zmiz1*-KO mice, we performed SOX2 and TBR2 immunostaining to quantify apical and intermediate progenitors, respectively, at embryonic day (E) 15.5 (Fig. 2A). We found a significant reduction in SOX2+ and TBR2+ progenitors in *Zmiz1*-KO compared to controls (Fig. 2B-E). To assess whether progenitor proliferative activity was impaired upon loss of *Zmiz1*, we performed BrdU birth-dating at E12.5, to target apical progenitors during production of deep-layer neurons, and at E14.5, to target intermediate progenitors during upper layer neuron production. We analyzed the number and distribution of BrdU+ neurons across the cortex at P7 (Fig. 2F, J). There is a significant decrease in BrdU+ neurons born at E12.5 in the deep layers (Fig. 2G-I), and in BrdU+ neurons born at E15.5 in upper layers in *Zmiz1*-KO compared to controls (Fig. 2K-M). These results indicate that neurons produced from apical and intermediate progenitors migrate appropriately to the layers corresponding to their birthdate, however, their production throughout neurogenesis is reduced upon *Zmiz1* mutation. Overall, these results indicate that the cortical microcephaly in *Zmiz1*-KO brains stemmed from reductions in cortical apical and intermediate progenitor populations due to their impaired proliferative abilities.

**Figure 2:**
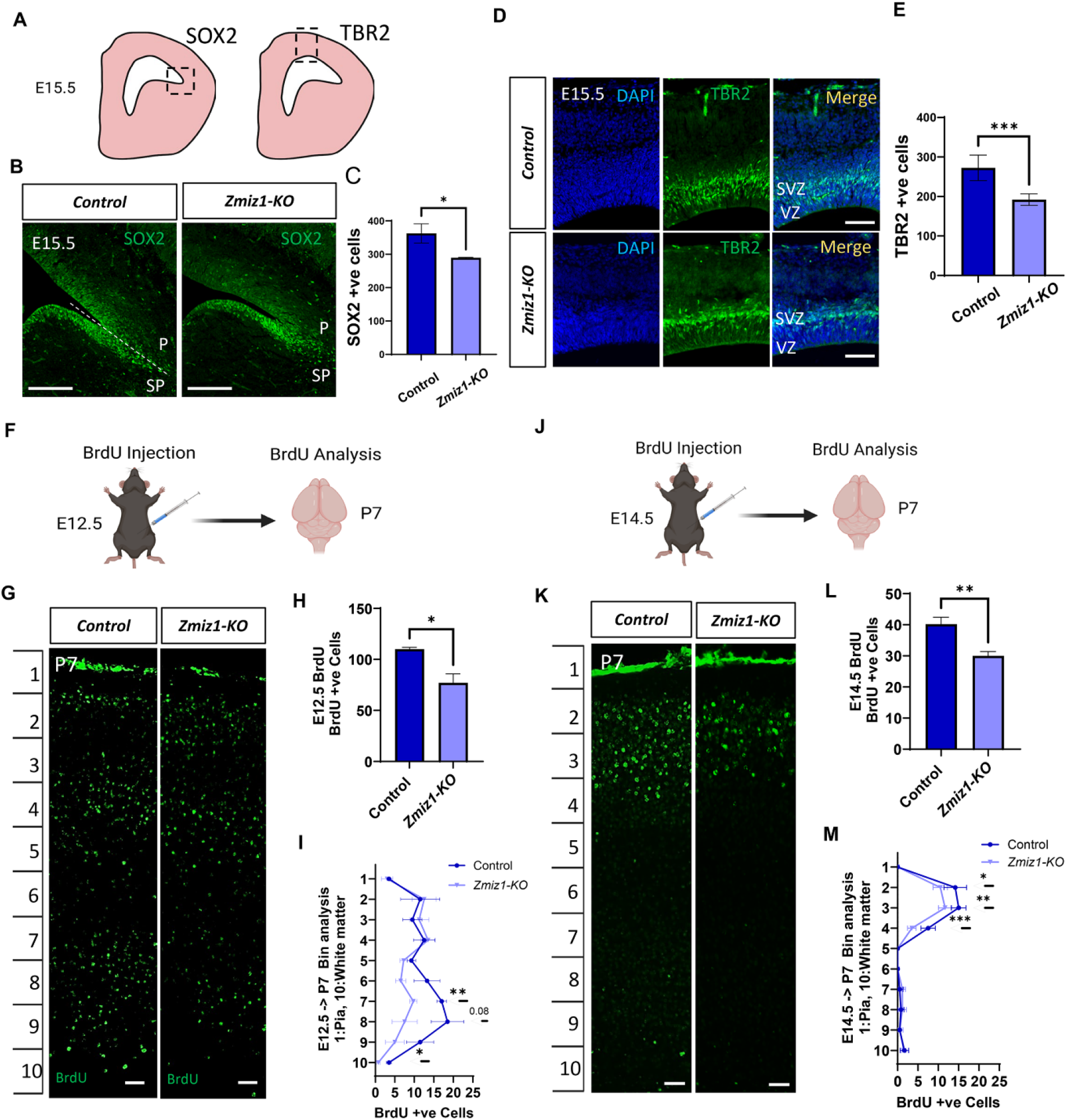
Loss of *Zmiz1* impairs cortical progenitor proliferation in the developing cortex. (A) Schematic of areas of E15.5 brains used for SOX2 and TBR2 quantification in (B) and (D). (B) Immunofluorescent antibody staining of SOX2 in brain sections show reduced SOX2+ progenitors in *Zmiz1*-KO cortex. (C) Quantification of SOX2+ cells in (B). (D) Immunofluorescent antibody staining of TBR2 in brain sections show reduced TBR2+ intermediate progenitors in *Zmiz1*-KO cortex. (E) Quantification of TBR2+ cells in (D). (F-I) BrdU-bith dating at E12.5. (F) Schematic of experimental approach, (G) BrdU+ neurons produced by progenitor at E12.5 in *Zmiz1*-KO and control cortices at P7, (H) quantification of BrdU+ cells in (G), and (I) analysis of BrdU+ neurons location across the depth of cortex represented in 10 equal bins for quantification. (J-M) BrdU-bith dating at E14.5. (J) Schematic of experimental approach, (K) BrdU+ neurons produced by progenitor at E14.5 in *Zmiz1*-KO and control cortices at P7, (L) Quantification of BrdU+ cells in (K), and (M) analysis of BrdU+ neurons location across the depth of cortex. Data are presented as mean ± SEM. * P <0.05, **P <0.01, ***P < 0.001, unpaired two-tailed t test. n = 3-4 per group. P –pallium, SP – subpallium, VZ – ventricular zone, SVZ – subventricular zone. Scale bars: 200 µm (B), 150 µm (D), 100 µm (G, K).

### Loss of *Zmiz1* results in corpus callosum dysgenesis and abnormal differentiation of layer 2/3 neurons

The strong reduction in midline length observed in *Zmiz1*-KO brains (Fig. 1A, C) suggests abnormal development of midline structures. The corpus callosum (CC) is a midline structure frequently affected in NDDs (19–23), particularly in patients with ID and/or ASD (19 23). Importantly, abnormalities in the CC and structures surrounding the midline have been reported in patients with ZMIZ1 mutation (9,24). Therefore, we investigated whether the CC and surrounding midline structures are affected in *Zmiz1*-KO mice.

First, we assessed the CC structure, length, and area using Nissl and gold chloride staining in adult mice. The CC area and length are significantly reduced in *Zmiz1*-KO compared to controls (Fig. 3A-E). The orientation of callosal fibers is also abnormal. We observed groups of fibers that were unable to cross the midline and followed abnormal trajectories extending into the septum or forming Probst bundles (Supplementary Fig. 3A-H). Dysgenesis of the CC could be due to abnormal development of callosal projection neurons, alterations in the midline structures guiding callosal axons across the midline (25,26) or both. We evaluated whether midline structures, such as the Glial wedge, Midline glial zipper (MGZ), Glial sling (GS), and Indusium griseum (IG) are abnormal and may contribute to the CC dysgenesis in *Zmiz1*-KO mice. We examined astroglia using GFAP immunolabeling at P7 and found a significant reduction of GFAP signal in *Zmiz1*-KO compared to controls. GFAP was particularly reduced in the MGZ and the GS, at the ventral boundary between the CC and septum (Fig. 3F-G). GFAP signal was distributed asymmetrically with respect to the midline (Fig. 3F-G), indicating abnormal location of astroglia in midline structures of *Zmiz1*-KO mice. The IG, at the dorsal border of the CC, was also affected in *Zmiz1*-KO (Supplementary Fig. 3A’, C’, E’, G’, I-N). Using TBR1 and CTIP2 as IG neuron makers, we revealed that IG neurons do not form a proper dorsal boundary separating CC axons and the interhemispheric fissure (Supplementary Fig. 3I-N). These results showed that midline structures developed abnormally and may contribute to CC dysgenesis in *Zmiz1*-KO mice. To directly assess the contribution of midline astroglia to the CC dysgenesis phenotype, we performed astrocyte-specific deletion of *Zmiz1* using GFAP-Cre. Astrocyte-specific *Zmiz1* deletion resulted in disorganization of astrocytes at midline, but not in dysgenesis or structural alterations in the CC (Fig. 3H-N). Together, these results strongly suggest that the callosal dysgenesis in *Zmiz1*-KO mice is primarily caused by the effect of *Zmiz1* deletion in neurons.

**Figure 3:**
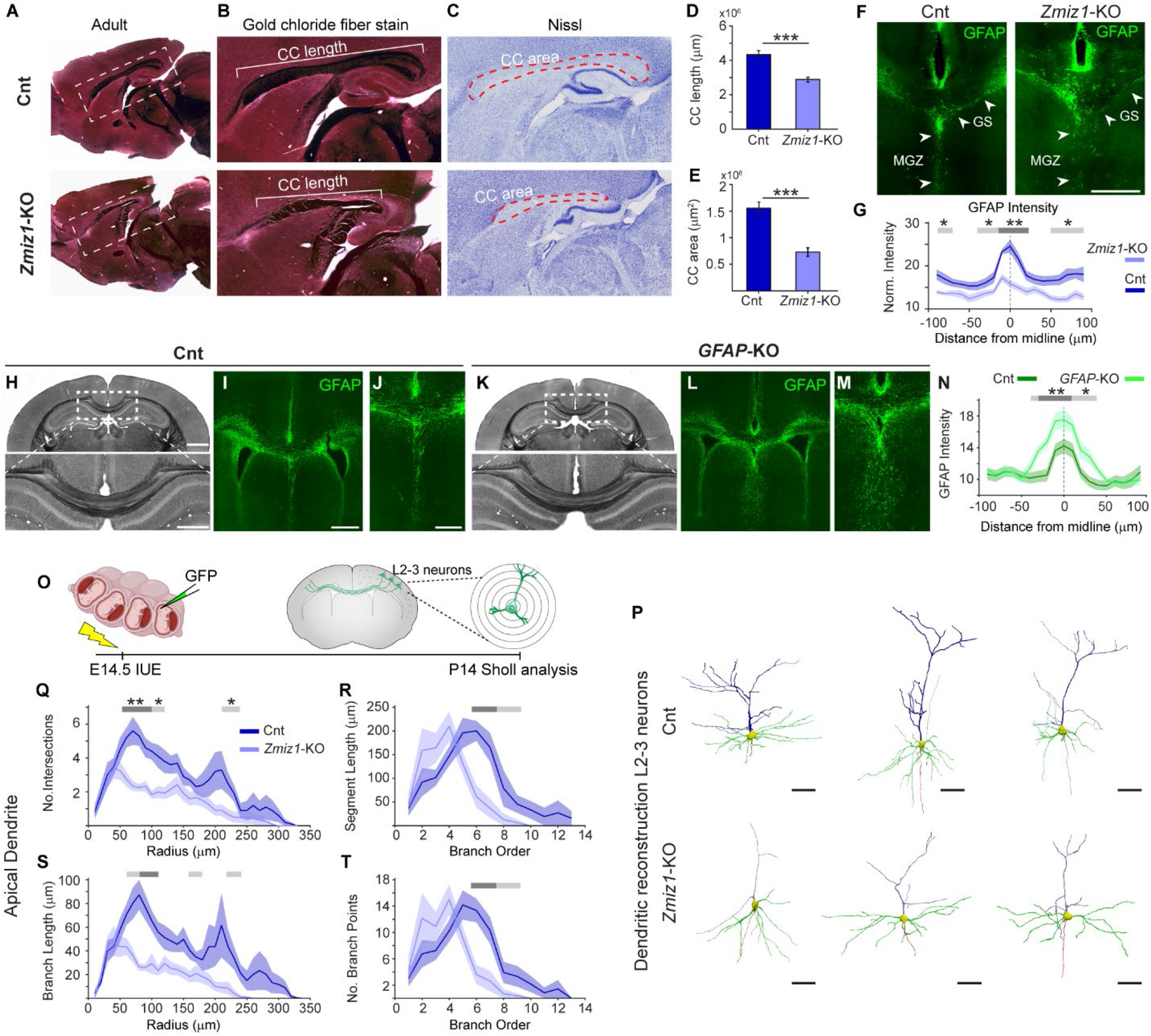
*Zmiz1* deficiency results in corpus callosum dysgenesis and abnormal differentiation of layer 2/3 neurons. (A-D) Morphological analysis of the corpus Callosum (CC). (A-B) Gold chloride staining for myelin axon bundle. Boxed areas in (A) are shown at higher magnification in (B), and Nissl-stained adjacent sections are shown in (C). (D, E) Quantification of CC area (D) and length (E). n = 10 per group. (F) GFAP staining showing abnormal midline astrocyte distribution at P7 in *Zmiz1*-KO brains. (G) GFAP intensity analysis in midline region shown in (F), n = 3 per group. (H, K) Gold chloride staining showing no callosal defects in *GFAP*-KO (*GFAP-Cre; Zmiz1*^*fl/fl*^). (I, J, L, M) GFAP staining showing astrocyte disorganization at the midline in *GFAP*-KO. (N) GFAP intensity plot for (J, M). (O) Schematics of experimental approach. (P) Representative reconstructed layer (L) 2/3 neurons at P14. n = 12 per group. (Q-T) Sholl analysis of Apical dendrites on reconstructed L2/3 neurons shows (Q) reduced number of intersection/arborization, (R) reduction of segment length in higher-order branches, (S) reduced length, and (T) reduction of higher-order branches in *Zmiz1*-KO L2/3 neurons. Data are presented as mean ± SEM. Two-tailed t-test (D, E), ANOVA-Tukey (G, N), * P <0.05, **P <0.01, ***P < 0.001, Scale bar: 1 mm (A-C), 100 µm (F, I), 1 mm (H, top), 500 µm (H, bottom), 50 µm (J, P).

We showed that cortical progenitors are affected, and fewer neurons are produced in *Zmiz1*-KO mice. However, whether and how loss of *Zmiz1* function alters neuron differentiation is still unclear. Since the majority of the CC is comprised of axons from L2/3 neurons (27), we focused on this population to investigate functions of *Zmiz1* in neuron differentiation. We first evaluated neurite outgrowth in differentiated Neuro-2a (N2a) cells in cultures treated with *Zmiz1* siRNA or control siRNA. *Zmiz1* knockdown resulted in significantly reduced neurite outgrowth compared to control siRNA-treated N2a (Supplementary Fig. 4A, B). Next, we examined the axonal and dendritic projections of cortical L2/3 neurons in vivo. L1CAM immunolabeling marks axonal callosal projections (28); we observed L1CAM projections extending abnormally in the CC and forming Probst bundles in *Zmiz1*-KO P7 mice (Supplementary Fig. 3B, D, F, H). Furthermore, L2/3 neurons were sparsely labeled to analyze dendritic arborizations. We performed *in utero* electroporation of a GFP-expressing construct in *Zmiz1*-KO and control embryos at E15.5 followed by dendritic reconstruction and Sholl analysis at P14 (Fig. 3O). We discovered a substantial disruption in the apical dendritic arbors of *Zmiz1*-KO L2/3 neurons, but not in the basal arbors (Fig. 3P-T, Supplementary Fig. 3O-R). The apical dendrite length, branching, and complexity are significantly reduced in *Zmiz1*-KO L2/3 neurons compared to controls (Fig. 3P-T). Examination of the branching patterns revealed that dendrite branch order and the length of the most distal segments were significantly reduced in apical arbors. This is indicative of a significant reduction of the apical tufts in *Zmiz1*-KO L2/3 neurons (Fig. 3R, T). Taken together, these data indicate that loss of *Zmiz1* leads to callosal dysgenesis, primarily due to abnormal differentiation of L2/3 neurons, which results in the aberrant formation of axonal projection and apical dendritic arbors.

### *Zmiz1* deficiency results in ASD-related behavioral phenotypes

To determine whether cortical deletion of *Zmiz1* in mice leads to behavioral alterations related to the developmental delays, ID, ASD, and ADHD present in human patients with ZMIZ1 mutations, we performed a comprehensive behavioral assessment of *Zmiz1*-KO mice. We examined behaviors associated with these disorders, including locomotor activity, anxiety-like behaviors, sensory gating, social interactions, and communication (Fig. 4, Supplementary Fig. 5). Since sex differences in these behaviors have been detected in mouse models of neurodevelopmental disorders (29–32), we assessed males and females independently.

**Figure 4:**
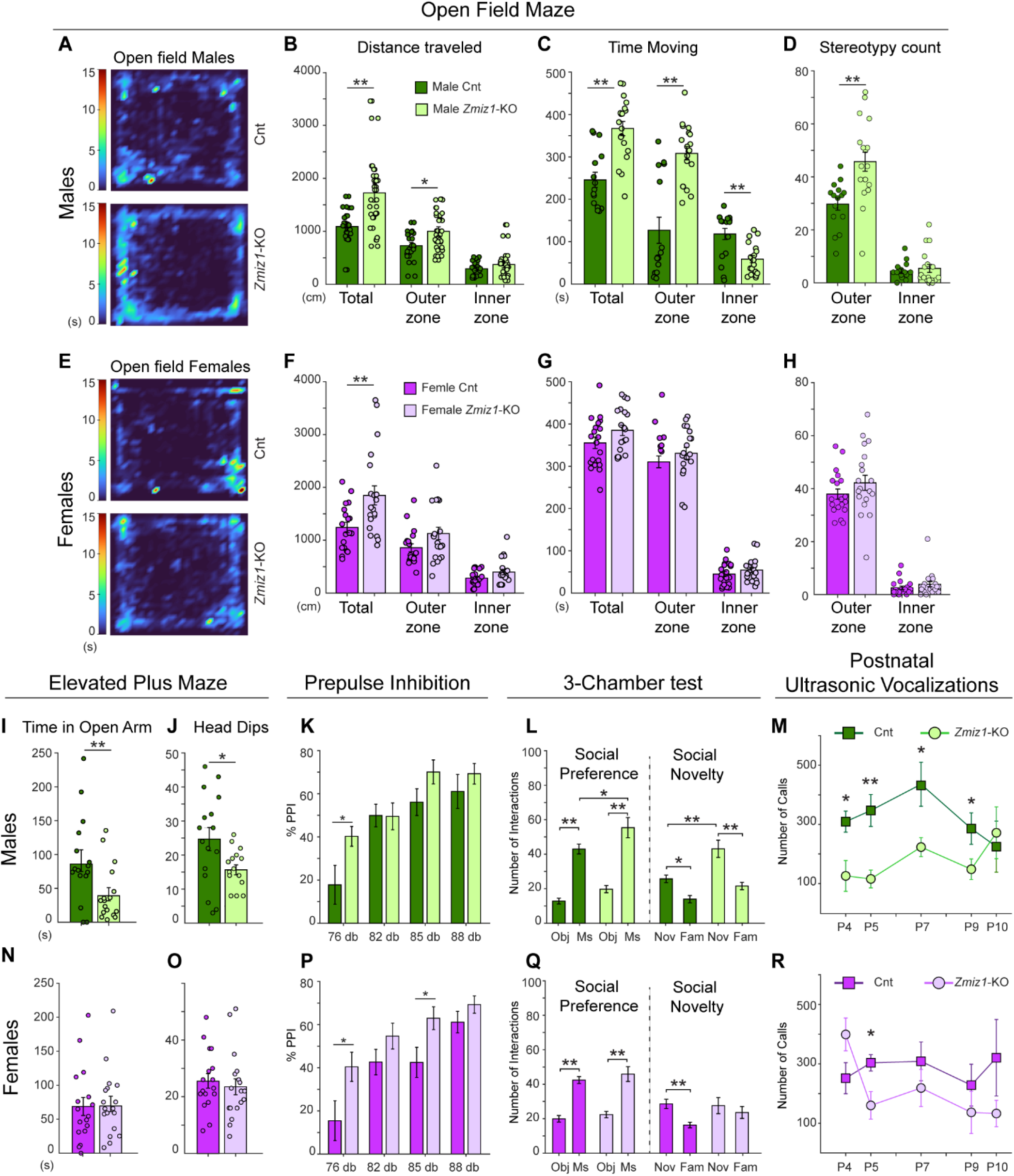
Loss of *Zmiz1* results in ASD-related behavioral phenotypes. (A, E) Representative heatmap plot showing locomotion of males (A) and females (E) in an open field test as a function of time. (B, F) Distance traveled by males (B) and females (F) in outer and inner zones of open field. (C, G) Time spent moving by males (C) and females (G) in outer and inner zones of open field. (D, H) Stereotypy activity count by males (D) and females (H) in outer and inner zones of the open field. (I, N) Time spent by males (I) and females (N) in the open arm of the elevated plus maze. (J, O) Number of head dips by males (J) and females (O) in elevated plus maze. (K, P) Percent of prepulse inhibition in males (K) and females (P) to tones of different intensities. (L, Q) Social preference and social novelty assessment of males (L) and females (Q) in three-chamber test. (M, R) Number of ultrasonic vocalizations (calls) in males (M) and females (R) pups at P4, P5, P7, P9, and P10. Data are presented as mean ± SEM. n = 12-18 per sex (control) and n = 12-18 per sex (*Zmiz1*-KO). * P <0.05, **P <0.01, calculated by unpaired two-tailed t-test, ANOVA-repeated measure (M, R).

In the open field test, we measured locomotor activity and anxiety, as it is a conflict test based on the opposing driver to explore and the fearful avoidance of exposed areas (107). *Zmiz1*-KO males and females traveled considerably more distance in the open field than control mice (Fig. 4A, B, E, F). However, only *Zmiz1*-KO males traveled significantly more distance in the outer zone compared to controls (Fig. 4B, F). Also, *Zmiz1*-KO males, but not females, spent substantially more time moving in the outer zone and less in the inner zone (Fig. 4C, G). These results indicate increased locomotor activity in *Zmiz1*-KO mice and suggest increased anxiety in males, as they tend to avoid exploration of the exposed inner zone. To further explore anxiety-related behaviors, we assessed stereotypies, including repetitive head bobbing and rearing in the open field arena. We found that *Zmiz1*-KO males exhibited significantly more episodes of stereotypy in the outer zone compared to controls, but not females (Fig. 4D, H). An independent assessment of anxiety-related behaviors using the elevated plus maze also revealed differences in male but not female *Zmiz1*-KO mice. Males spent less time exploring the open arms of the elevated plus maze and showed decreased occurrences of exploratory behaviors such as head dips (Fig. 4I, J, N, O), both indicative of increased anxiety.

Next, we assessed sensory gating using the acoustic startle/prepulse inhibition (PPI) test. Males showed significantly less PPI in response to lower prepulse levels, and females showed significantly less PPI with lower and intermediate prepulse levels (Fig. 4K, P). These results suggest a mild impairment in sensory gating in *Zmiz1*-KO males and females compared to controls. In addition, we evaluated working memory using T-maze. We found no difference in performance compared to controls in either *Zmiz1*-KO males or females (Supplementary Fig. 5A).

Social deficits are a common phenotype of many neurodevelopmental disorders and present in patients with ZMIZ1 mutations (7,9,10). We evaluated social preference and social novelty in *Zmiz1*-KO mice using the three-chamber test. Both male and female *Zmiz1*-KO mice and controls, interacted more with a conspecific than with an object, demonstrating normal social preference (Fig. 4L, Q). However, *Zmiz1*-KO males exhibit higher interaction with the novel conspecific than control males (Fig. 4L). When tested for social novelty preference, *Zmiz1*-KO males interacted more with novel mice than with familiar mice and showed more interaction with novel mice than control males (Fig. 4L). In contrast, *Zmiz1*-KO females showed no preference for the novel compared to the familiar conspecifics, unlike control females (Fig. 4Q). These results indicate that *Zmiz1*-KO males and females exhibit altered social interactions, however, while males showed increased social novelty preference compared to controls, females exhibit no social novelty preference.

Delayed speech onset and deficits in communication are universal in patients with ZMIZ1 variants (7,9,10). To investigate whether *Zmiz1*-KO mice present early communication deficits, we examined neonatal ultrasonic vocalization (USV) emitted by pups upon isolation from the mother at P4, P5, P7, P9, P10. Overall, *Zmiz1*-KO pups emitted fewer USV than controls, suggesting communication deficits in their early life (Fig. 4M, R). More specifically, *Zmiz1*-KO males showed significant reduction in the numbers of calls at P4, P5, P7, and P9 compared to controls, while *Zmiz1*-KO females only showed significant decrease at P5 (Fig. 4M, R, Supplementary Fig. 5B). Moreover, we analyzed the call types or syllables and found that *Zmiz1*-KO males produce significantly less variety of syllables compared to male controls and females (Supplementary Fig. 5C-F). These alterations are likely independent of altered respiratory function and their physical ability to make calls, since the mutation only affects the forebrain, there are no significant differences in the pup’s weight, and other USV parameters are normal (Supplementary Fig. 5G-J). Collectively, *Zmiz1*-KO mice exhibit behavioral alterations with different manifestations in males and females and are present from early postnatal stages to adulthood. Importantly, these deficits in communication, motor activity, anxiety, sensory gating, and social interactions recapitulate aspects of human neurodevelopmental phenotypes in ZMIZ1 human patients.

### *Zmiz1* loss alters expression of genes associated with neurogenesis, neuron differentiation, and synaptic transmission

To better understand the molecular role of *Zmiz1* in the cortex, we performed transcriptome analysis. Given the functions of *Zmiz1* in transcriptional regulation, RNA sequencing (RNA-seq) experiments were utilized to examine genome-wide gene expression changes induced by *Zmiz1* mutation in cortical progenitors, immature neurons, and mature neurons. RNA-seq was conducted on isolated E15.5 progenitors, and on P1 and P14 neurons from controls and *Zmiz1*-KO cortices (Fig. 5A, E, I). Analysis of differentially expressed genes (DEGs) analysis revealed 67 upregulated and 194 downregulated genes in E15.5 *Zmiz1*-KO progenitors (E15.5 DEGs), 124 upregulated and 167 downregulated genes in P1 *Zmiz1*-KO neurons (P1 DEGs), and 749 upregulated and 1367 downregulated genes at P14 *Zmiz1*-KO neurons (P14 DEGs) (Supplementary Fig. 6A-C). DEGs were further evaluated for functional enrichment. Gene ontology (GO) for biological processes analysis revealed neuron development as broadly affected biological process in all three populations. However, biological processes such as axon guidance, axonogenesis neuron projection guidance/development, neurogenesis, and neuron differentiation were also affected in E15.5 progenitors and P1 neurons (Fig. 5B, F, J). Interestingly, P14 DEGs were heavily associated with synaptic transmission and signaling (Fig. 5J). KEGG analysis shows various affected processes, such as synaptic vesicle cycle, axon guidance, glutamatergic synapse, nicotine addiction, and GABAergic synapse at the different stages (Supplementary Fig. 7A, D, G). Reactome pathway shows EPH-ephrin signaling, L1CAM interactions, SEMA3A signaling and interaction, protein-protein interactions at synapses, and synaptic signal transmission as affected processes (Supplementary Fig. 7B, E, H), and Jensen Compartment subcellular location database shows dendrite/dendritic tree, axon/axon collateral, and synapse/pre-synapse/post-synapse as the compartments preferentially associated with P14 DEGs (Supplementary Fig. 7C, F, I).

**Figure 5:**
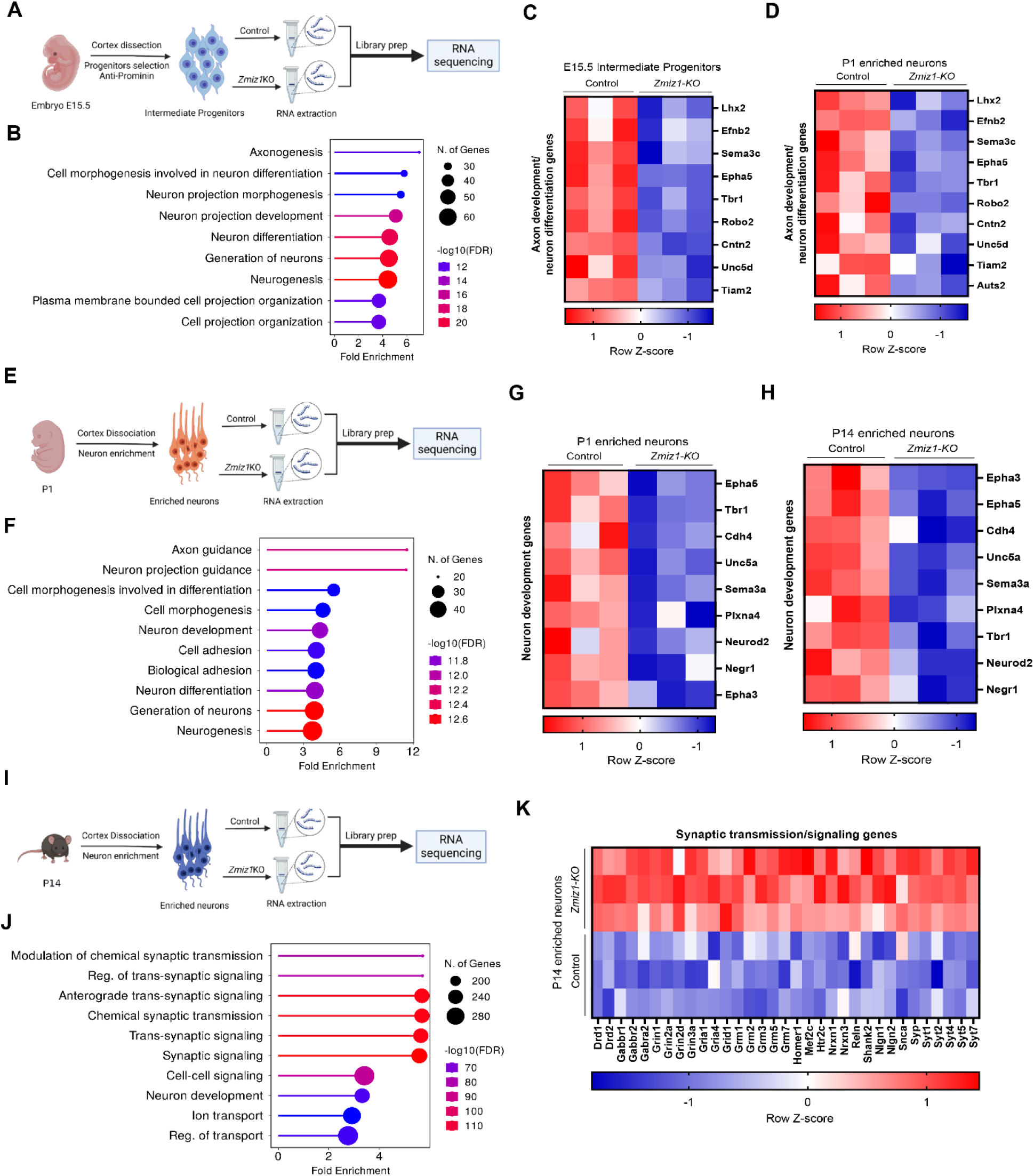
*Zmiz1* loss alters expression of genes associated with neurogenesis, neuron differentiation, and synaptic transmission. (A, E, I) Schematic depicting intermediate progenitor isolation from E15.5 cortex (A) and neuron enrichment from P1 (E) and P14 cortex (I). (B, F, J) Gene ontology (GO) analysis for biological processes on E15.5 DEGs (B), P1 DEGs (F), and P14 DEGs (J). (C, D) Heatmap for E15.5 (C) and P1 (D) axon development/neuron differentiation DEGs. (G, H) Heatmap for P1 enriched neuron (G) and P14 enriched neuron (H) development DEGs. (K) Heatmap for P14 enriched neurons synaptic transmission/signaling DEGs.

We found comparable changes in expression at E15.5 and P1 in axon development/neuron differentiation DEGs. Notably, *Lhx2, Efnb2, Sema3c, Epha5, Tbr1, Robo2, Cntn2, Unc5d*, and *Tiam2* were axon development/neuron differentiation DEGs downregulated in both E15.5 progenitors and P1 neurons in *Zmiz1*-KO cortex, suggesting that *Zmiz1* may regulate these genes throughout early developmental stages (Fig. 5C, D). Similarly, neuron development DEGs such as *Epha5, Tbr1, Cdh4, Unc5a, Sema3a, Plxna4, Neurod2*, and *Negr1* were downregulated in both P1 and P14 neurons (Fig. 5G, H). Interestingly, P14 DEGs were highly enriched in synaptic transmission/signaling genes including *Drd1/2, Gabbr1/2, Gabbra2, Grin1/2a/2d/3a, Gria1/4, Grid1, Grm1/2/3/5/6, Homer1, Mef2c, Htr2c, Nrxn1/3, Reln, Shank2, Nlgn1/2, Snca, Syp*, and *Syt1/2/4/5/7* (Fig. 5K). Overall, transcriptomic analysis in progenitors, immature neurons, and mature neurons suggests that *Zmiz1* regulates expression of genes controlling neuron development and differentiation early in development, and synaptic signaling genes later in development.

### Abnormal regulation of synaptic-related and ASD/ID-risk genes in *Zmiz1*-KO cortex

To investigate the link between *Zmiz1*-KO DEGs and *Zmiz1*-associated neurodevelopmental disorders, we performed correlation analysis between DEGs and ASD (SFARI) and ID (33) risk gene databases (Supplementary Fig. 8A). A significant number of DEGs caused by *Zmiz1* mutation were disease risk factors for ASD and ID. Seven ID-genes and 31 ASD-genes mapped to E15.5 DEGs, 13 ID-genes and 51 ASD-genes mapped to P1 DEGs, and 90 ID-genes and 244 ASD-genes mapped to P14 DEGs (Supplementary Fig. 8B-D). Five genes, *Auts2, Plcb1, Satb2, Syp*, and *Tbr1*, were mapped to all three datasets (SFARI, ID risk, and *Zmiz1*-KO DEGs) at E15.5 (Supplementary Fig. 8B). Four genes, *Auts2, Foxp1, Lama1*, and *Tbr1*, mapped to the three datasets at P1 (Supplementary Fig. 8C), and 42 genes, including *Cntnap2, Gria3, Grin1, Grin21, Nrxn1, Reln, Satb2, Shank2, Syngap1*, and *Tbr1*, mapped to the three datasets at P14 (Supplementary Fig. 8D).

Gene Ontology analyses revealed a strong involvement of *Zmiz1*-KO DEGs in neuron development and differentiation, and synaptic transmission processes. Neuron differentiation and synaptic communication are thought to produce alterations in excitatory/inhibitory balance leading to circuit dysfunction in ASD and other NDDs (34–37). To investigate whether *Zmiz1* loss preferentially affects excitatory vs inhibitory neurons, we mapped DEGs to markers of specific cell types, either excitatory or inhibitory (38). We found that *Zmiz1*-KO DEGs were genes mainly expressed in excitatory neurons (Supplementary Fig. 8E). Additionally, we mapped the DEGs to the excitatory vs inhibitory synaptic cleft proteome (39). We found 10 E15.5 DEGs, 23 P1 DEGs, and 94 P14 DEGs are proteins found in excitatory synapses (including *Gria1/2/4, Grin1/2b, Nrxn1/2, Shisa7*). In contrast, K C at al, 2024 [preprint] 12 of 27 3 E15.5 DEGs, 3 P1 DEGs, 25 P14 DEGs correspond to inhibitory synaptic proteins (including *Gabra1/2/3, Cdh10, Gabrb1/2/3, Nrxn3, Slc6a1, Gabbr1, Reln*) (Supplementary Fig. 8F). Also, we found 2 E15.5 DEGs, 1 P1 DEG, and 9 P14 DEGs genes correspond to proteins found in both excitatory and inhibitory synapses (including *Cntnap1, Nrxn2, EphB6, Tenm2, Nlgn1*) (Supplementary Fig. 8F).

Given the enrichment of GO pathways involved in synaptic function, we further compared the *Zmiz1*-KO DEGs with the synaptic gene ontology database SynGO (40). Forty-two E15.5 DEGs, 62 P1 DEGs, and 437 P14 DEGs were identified as synaptic genes (Supplementary Fig. 8G-I). GO analysis of DEGs-synaptic genes showed enrichment of biological processes such as synapse organization, neurotransmitter transport, and regulation of neurotransmitter levels at E15.5 (Supplementary Fig. 8J); nicotine addiction, long-term depression, long-term potentiation, and glutamatergic synapse at P1 (Supplementary Fig. 8K); and nicotine addiction, long-term potentiation, synaptic vesicle cycle, and glutamatergic synapse at P14 (Supplementary Fig. 8L). These data indicate that *Zmiz1* plays a crucial role in directly or indirectly regulating transcription of ID/ASD risk genes involved in synaptic function and neuron development.

### *Zmiz1* regulates expression *of Auts2, Efnb2, Lhx2, EphA5*, and *Unc5d*

To further investigate the molecular mechanism and downstream targets regulated by *Zmiz1*, we performed ChIP-sequencing (ChIP-seq). HA-tagged ZMIZ1 (*Zmiz1*-HA) was overexpressed in N2a cells followed by ChIP-seq analysis (Fig. 6A, Supplementary Fig. 9). ZMIZ1 ChIP-seq revealed thousands of peaks and GTGTATGTGTGT as the top *Zmiz1* motif (Supplementary Fig. 10, Supplementary Fig. 11). We integrated ZMIZ1 ChIP-seq and RNA-seq data to reveal directly regulated targets. When comparing the ChIP peaks with the groups of E15.5 and P1 DEGs involved in axon guidance/neuron differentiation we found 6 genes overlapping in these 3 datasets: *Auts2, Lhx2, Efnb2, Epha5, Unc5d*, and *Slit2* (Fig. 6B). All these genes in the RNA-seq datasets from *Zmiz1*-KO cortices were found to be downregulated in the E15.5 progenitors and P1 neurons RNA-seq datasets from *Zmiz1*-KO cortex, except for Slit2 which is upregulated at E15.5 and downregulated at P1 (Supplementary Fig. 11B).

**Figure 6:**
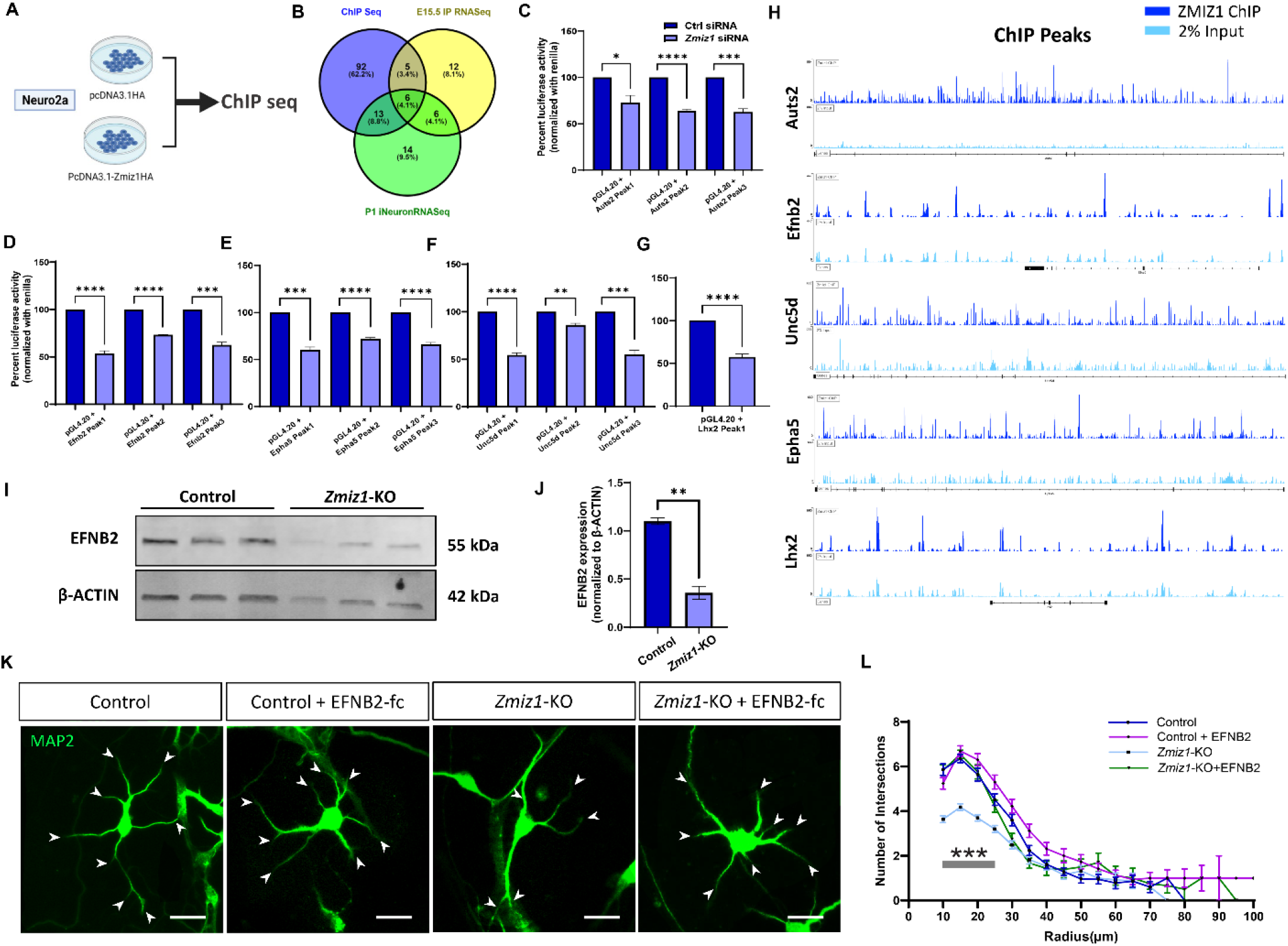
*Zmiz1* regulates expression of Efnb2, *Auts2, Lhx2, Unc5 and Epha5*, and *Efnb2* treatment rescues neurite branching defects in *Zmiz1*-KO cortical neurons in vitro. (A) Schematic of ZMIZ1-HA ChIP-sequencing. (B) Venn diagram showing overlapping axon guidance and differentiation genes from ChIPseq peaks, E15.5 intermediate progenitors DEGs, and P1 enriched neurons DEGs. (C-G) Percent luciferase activity for *Auts2* (C), *EfnB2* (D), *EphA5* (E), *Unc5d* (F), and *Lhx2* (G) ChiP-seq peaks is decreased upon *Zmiz1* knockdown in N2a cells. (H) ChIP-seq peak genomic plot for *Auts2, Efnb2, Unc5d, Epha5*, and *Lhx2*. (I) Western blot showing reduction in EFNB2 protein in *Zmiz1*-KO cortex. (J) Densitometric quantification of (I). (K) P1 control and *Zmiz1*-KO cortical neuron cultures fluorescently immunolabeled for MAP2 (green) showing rescue of neurites branching in *Zmiz1*-KO neurons upon EFNB2 treatment. Arrowheads indicate neurites. (L) Sholl analysis on reconstructed P1 cortical neurons (n = 50 per group, asterisks indicate P <0.001 for the three comparisons between the *Zmiz1*-KO group with the Control, Control + EFNB2, and *Zmiz1*-KO + EFNB2 groups). Data are presented as mean ± SEM. * P <0.05, **P <0.01, ***P < 0.001, unpaired two-tailed t-test. Scale bar: 50 µm (J).

To further confirm that ZMIZ1 regulates transcription of these potential downstream targets, we cloned ZMIZ1 ChIP peaks found in *Auts2, Lhx2, Efnb2, Epha5*, and *Unc5d* into a luciferase activity reporter vector and performed luciferase assay. We found a significant decrease in luciferase expression from *Auts2, Lhx2, Efnb2, Epha5*, and *Unc5d* ChIP peaks upon *Zmiz1* knockdown in N2a compared to siRNA control treated cells (Fig. 6C-G). In these genes, peaks bound by ZMIZ1 were found in intronic or intergenic loci (Fig. 6H, Supplementary Fig. 10, Supplementary Fig. 11B). These results suggest that ZMIZ1 binding directly regulates transcription of these genes. Overall, RNA-seq, ZMIZ1 ChIP-seq, and luciferase data implicate direct regulation of *Auts2, Efnb2, Lhx2, Epha5*, and *Unc5d* gene expression in the cortex by ZMIZ1.

### EFNB2 treatment rescues neurite branching defects observed in *Zmiz1*-KO cortical neurons

Eph-ephrins are receptor-ligand cell surface proteins that play essential roles in axon guidance, neurite outgrowth, and neuronal connectivity (41–44), and importantly, EphrinB2 (Efnb2) is critical for the development of the CC (44,45). We showed that Efnb2 is a target of *Zmiz1*, and its mRNA expression is significantly decreased in *Zmiz1*-KO cortex at E15.5 and P1. Given the callosal dysgenesis and abnormal dendritic development present in *Zmiz1*-KO mice, we investigated the hypothesis that dysregulation of the Efnb2 pathway may contribute to these phenotypes.

First, we assessed EFNB2 protein levels and found it is significantly decreased in the *Zmiz1*-KO cortex (Fig. 6I, J). Next, we focused on investigating *Efnb2* function in neurite outgrowth using *Zmiz1*-KO and control cortical neuron cultures. To assess neurite growth, we cultured neurons 5 days in vitro (DIV), immunolabeled for MAP2, and performed Sholl analysis on reconstructed neurons. We observed a significant decrease in neurite branching in *Zmiz1*-KO neurons compared to wildtype control neurons (Fig. 6 K, L). Next, we assessed the effect of *Efnb2* treatment on neurite growth. We applied EFNB2-fc recombinant protein to directly activate the Eph-ephrin activity in control and *Zmiz1*-KO cortical neurons at 2 DIV and assessed the neurite growth at 5 DIV. We compared neurons from Control, Control + EFNB2-fc, *Zmiz1*-KO, and *Zmiz1*-KO + EFNB2-fc groups. *Zmiz1*-KO neurons showed significantly reduced branching compared to controls (Fig. 6 K, L). While neurite branching was not significantly different in EFNB2-fc treated control neurons compared to non-treated controls, the neurite branching defect in *Zmiz1*-KO neurons was statistically rescued to control numbers upon EFNB2-fc treatment (Fig. 6 K, L; P <0.001 for comparison of *Zmiz1*-KO group with either Control, Control + EFNB2-fc, *Zmiz1*-KO + EFNB2-fc). Together, our data demonstrate that Efnb2 is a downstream target of *Zmiz1*, that dysregulation of the Efnb2 pathway contributes to *Zmiz1*-KO phenotype, and that reactivation of the Efnb2 pathway rescue neurite branching defect in *Zmiz1*-KO neurons in vitro.

## DISCUSSION

ZMIZ1 mutations are present in patients with a range of NDDs, including ASD and ADHD, and have recently been identified as the cause of a neurodevelopmental syndrome with microcephaly, ID, and craniofacial abnormalities as core clinical features (46–53). ZMIZ1 has also been classified as a strong candidate ASD risk gene by SFARI. Despite its strong association with NDDs, the functional roles of ZMIZ1 in brain development are completely unknown. Understanding how ZMIZ1 contributes to NDDs will require functional mechanistic studies that are not feasible in humans. To address this challenge, we generated a forebrain-specific *Zmiz1* mutant model that recapitulates phenotypes observed in the human ZMIZ1 neurodevelopmental syndrome, and other NDDs, including cortical microcephaly, callosal dysgenesis, and ASD-related behaviors (7–11). Our studies reveal that *Zmiz1* regulates cortical neurogenesis and different aspects of neuron differentiation, such as axon and dendritic development, and synaptic transmission. Thus, this work lays the foundation for further preclinical and mechanistic studies on ZMIZ1 function.

In the cortex, *Zmiz1* is expressed in progenitors and postmitotic neurons (16). This study shows that *Zmiz1* regulates neurogenesis and neuron differentiation. DEGs associated with loss of *Zmiz1* in E15.5 progenitors and P1 immature neurons overrepresented biological processes such as neurogenesis, neuron development, and neuron differentiation. Importantly, we showed a significant reduction in apical and intermediate progenitors and neurons produced from them. This likely resulted in the thinner and smaller cortices of *Zmiz1*-KO brains. Therefore, these findings are consistent with the observation of microcephaly in most ZMIZ1 mutant patients, and the presence of thinner cortex in patients with specific variants (8,9), which highlights the validity of this mouse model.

We revealed that *Zmiz1* regulates differentiation of callosal projection neurons. *Zmiz1* has been identified as an early developmental gene enriched in callosal and corticothalamic projection neurons (15), however, its functions in these neuron subtypes had not been previously investigated. Our studies reveal that *Zmiz1* regulates differentiation of callosal projection neurons. Callosal neurons predominate in cortical upper layers. They are critical for inter-hemispheric communication through the CC, are involved in high-level cognitive functions, and their abnormal development has been linked to neurologic disorders such as ASD (27,54,55). Importantly, single nuclei RNA sequencing on cortices from autistic patients revealed disruption in the development and synaptic signaling of upper-layer excitatory neurons (38). Given their functional importance and contribution to ASD pathology, we focused on investigating the functions of *Zmiz1* in the differentiation of L2/3 callosal neurons. We showed that *Zmiz1* regulates axonal extension through the CC. Moreover, our differential expression analysis at E15.5 and P1 revealed that axonogenesis and axon guidance are processes highly impacted by *Zmiz1* deletion. Subsequently, these processes may be important for more focused, future investigation since impairment of the CC is one of the most consistent brain phenotypes seen in autistic patients (56,57). Reduction in the connectivity between the cerebral hemispheres is common in ASD and ID, it is linked to increased excitability in local cortical circuits (58), and contributes to cognitive, social, and communication deficits (20,59–61). For these reasons, our findings demonstrating abnormal callosal neuron differentiation, corpus callosum dysgenesis, and social and communication deficits in *Zmiz1*-KO mice are clinically relevant because they are consistent with the association of these anatomical and behavioral phenotypes in humans.

In addition to callosal dysgenesis, we found alterations in the apical dendrites of L2/3 neurons in the *Zmiz1*-KO mice. Alterations of dendritic morphology have been observed in individuals with ASD (62,63). L2/3 neuron apical dendrites play a critical role in supralinear dendritic integration and subsequently synaptic transmission and connectivity (64,65). Impairments in the L2/3 apical dendrites may result in alterations in excitation and inhibition balance, as observed in ASD and numerous neurological disorders (66), and may contribute to the behavioral deficits observed in the *Zmiz1*-KO mice. Interestingly, *Zmiz1* is a co-activator of *p53* (47) and *Notch1* (67–69), which are critical regulators of dendritic development and cortical circuitry. Whether these pathways contribute to abnormal dendritic morphology in L2/3 neurons, and potentially in other neuron types in *Zmiz1* mutants, will require further investigation.

Our investigation of molecular pathways downstream of *Zmiz1* identified targets highly relevant to NDDs. *Zmiz1* targets high-confidence autism risk genes linked to ID such as *Lhx2* and *Auts2* (70–76). Lhx2 controls cortical neurogenesis (72–75), while *Auts2* regulates neurite outgrowth and spines (71,77). Moreover, we identified the Eph-ephrin signaling pathway as a target of *Zmiz1*. Eph-ephrin signaling is critical for cortical neurogenesis, synaptic development, axon guidance and CC development (78–81). Specifically, *EfnB2* and *EphA5* are transcriptionally regulated by *Zmiz1. EphA5* and *EfnB2* receptor, *EphB2*, are critical for NMDAR interaction and synaptic function (81–85). Although we have not evaluated synaptic functions in *Zmiz1*-KO, we found a significant number of DEGs associated with synaptic transmission, particularly at P14. Interestingly, *Zmiz1* mRNA is enriched in neuropil compared to somata, and is present in extra-nuclear cytoplasmic compartment (14,86). This suggests that *Zmiz1* may function in synaptic transmission, perhaps via regulation of Eph-ephrin pathway.

Importantly, *EfnB2* regulates neurite outgrowth in cortical neurons (87) and we show that *Zmiz1*-KO neurons in culture have impairments in neurite growth and branching that can be reverted by EFNB2 treatment. Future studies will determine whether *Zmiz1* plays additional functions in regulating synaptic transmission, whether Eph-ephrin signaling mediates these potential synaptic functions, and whether Eph-ephrin signaling can potentially ameliorate phenotypes produced by *Zmiz1* dysfunction.

Other interesting *Zmiz1* targets associated with NDDs and classified as ASD risk genes are FOXP2 and TBR1. *Zmiz1* was identified as an early developmental gene enriched in callosal and corticothalamic neurons (15) and corticothalamic neurons strongly express TBR1 and FOXP2 (88–90), which regulate cortical development, neurite growth, and synaptic transmission, and are associated with ASD and NDDs (91–94). Corticothalamic neurons are important for the modulation of sensory information processing (95,96). Their dysfunction contributes to sensory gating impairments and social behavioral alterations in NDDs (97). Though we focused on L2/3 callosal neurons in this study, *Zmiz1* may regulate differentiation and synaptic transmission in other neuron types and circuits relevant to NDDs. Understanding *Zmiz1* functions in specific cortical neuron subtypes such as corticothalamic neurons requires future exploration.

In conclusion, our study demonstrates that *Zmiz1* is crucial for cortical development, and its loss of function results in cortical developmental defects and behavioral deficits that resemble human conditions with ZMIZ1 mutations. We also show that *Zmiz1* controls expression of key cortical development and NDD risk genes. Overall, our findings in *Zmiz1* mutant mice provide insight into the role of *Zmiz1* in cortical development, the molecular underpinnings of ZMIZ1 syndrome and will contribute to understanding ASD and NDD pathophysiology.

## ACKNOWLEDGMENT

We thank Dr. Jovanny Zabaleta and Dr. Jone Garai at The Louisiana Cancer Research Consortium (LCRC) Translational Genomics Core Center for their continued support of our sequencing experiments.

## Author Contributions

RKC, NRP, AT, SMM, and MJG developed and designed the experiments. RKC, AT, NRP, AST, AS, and MGL performed experiments. RKC, AT, and NRP analyzed data. AT, IK, MA, and VM analyzed behavioral data with guidance from MJG. RKC and MJG wrote the manuscript.

## SUPPLEMENTARY MATERIAL

Supplementary Figures S1-S11

## MATERIAL AND METHODS

### Mouse line and Breeding

All animal experiments were performed following Tulane University Institutional Animal Care and Use Committee (IACUC) policy. *Zmiz1*^*fl/fl*^ (98), *Emx1-Cre* (99) (Stock No. 005628, B6.129S2-Emx1tm1(cre)Krj/J), and GFAP-Cre (Stock No. 024098) were bred to conditionally ablate *Zmiz1* in the Emx1 lineage or GFAP lineage. All experiments with *Zmiz1*^*fl/fl*^*;Emx1-Cre* and *Zmiz1*^*fl/fl*^*;GFAP-Cre* were controlled with *Zmiz1*^*fl/fl*^ mice. For embryonic (E) studies and *in utero* electroporation, E0.5 was designated to the day when a vaginal plug was found. For postnatal studies, postnatal day (P) 0 was assigned to the day when a mouse was born.

### Immunohistochemistry

Immunohistochemistry was performed as previously described (100). Briefly, mice were transcardially perfused with ice-cold PBS and 4% PFA. Brains were dissected and post-fixed with 4% PFA at 4°C overnight. The brain was sectioned coronally on a vibratome (Leica). Sections were blocked using Cas block (ThermoFisher, 008120) and incubated with primary antibodies at 4°C overnight followed by 1-hour secondary antibody incubation the next day. For DAPI staining, tissue was mounted in DAPI Fluoromount-G (SouthernBiotech, 0100-20). Antibodies used include ZMIZ1 (Cell Signaling, 89500S), TBR2 (Ebiosciences 14-4875-82, 1:200), SOX2 (EMD Millipore, AB5603, 1:200), BRDU (Novus Biologicals, NBP2-14890, 1:200, or Abcam, ab142567), CUX1 (ProteinTech 11733-1-AP, 1:300), CTIP2 (Abcam, 18465, 1:500), TBR1 (Abcam 31940, 1:1000), NEUN (Sigma-Millipore MAB377, 1:500), GFP (Aves Labs, GFP-1020, 1:500), GFAP (Aves Labs, GFAP, 1:500), MAP2 (Abcam, ab5392, 1:5000), Phalloidin (Invitrogen, A12379, 1:500), L1CAM (Sigma-Millipore MAB5272, 1:500). Secondary antibodies include donkey anti-Rabbit 488 (Invitrogen, A21206), donkey anti-Rabbit 555 (Invitrogen, A31572), donkey anti-Rabbit 633 (Invitrogen, A21070), goat anti-Rat 555 (Invitrogen, A31572), goat anti-chicken 488 (Invitrogen, A11039), donkey anti-mouse 555 (Invitrogen, A31570), donkey anti-mouse 488 (Invitrogen, A11001).

### BrdU birth dating

Timed pregnancies were carried out and bromodeoxyuridine (BrdU) (100 mg/kg in PBS) was injected intraperitoneally at Embryonic day (E) 12.5 and 14.5. Postnatal day (P) 7 mice were perfused. P7 brains were then collected, sectioned, and processed for BrdU immunohistochemistry. (details in immunohistochemistry section)

### Gold chloride staining

Gold chloride staining was performed as previously described (101). Briefly, brain sections were washed with PBS and mounted on Superfrost Plus slides. Slides were incubated in 0.2% gold chloride solution at 37 °C for 30 - 90 minutes hours. Slides were rinsed with deionized water, followed by dehydration in ethanol, cleared in xylene, and mounted with DPX mounting medium (Sigma-Aldrich).

### Nissl Staining

Nissl staining was performed as previously described (102). Briefly, brain sections were washed with PBS and mounted on Superfrost Plus gelatin-coated slides. Sections were dehydrated and cleared in xylene followed by rehydration. Slides were incubated with prewarmed 0.5% Cresyl violet solution for 3-5 minutes followed by 50% ethanol and 0.1% acetic acid. The sections were dehydrated, and cover slipped using DPX mounting medium.

### *In utero* electroporation and neuron reconstruction

*In utero* electroporation of p-CMV/b-actin-GFP was performed as previously described (15). Briefly, constructs were electroporated in E14.5 embryos using a 30mV square pulse for 50ms. GFP expression was amplified using anti-GFP antibody and analyzed at P14. Cortical layer 2/3 neurons were imaged at 60X with 0.5 μm steps using a Nikon C2 confocal microscope. Neuron reconstruction and Sholl analysis were performed using Neurolucida 360 (MBF Bioscience).

### N2a differentiation

Neuro-2a cells (N2a) (ATCC, CCL-131) were cultured in DMEM-high glucose (Cytiva, SH30022) supplemented with 10 % FBS (Cytiva, SH3008803) and 1% Penicillin-Streptomycin (Thermo Fisher, 15070063) at 37 °C and 5% CO2. N2a cells were transfected with a pool of control non-targeting siRNA (Horizon, D-001810-10-05) or ZMIZ1 siRNA (Horizon, L-007034-00-0005) for 48 hours using Lipofectamine 3000 (Thermo Fisher, L3000015). The final concentration of siRNA solution was 200 nM. N2a cells were serum starved for two days and then stained with MAP2 (Abcam, ab5392, 1:5000) and Phalloidin (Thermofisher Scientific, A12379, 1:1000).

### Neuron culture

Cortices were dissected from P1 mice brains and dissociated using Papain Dissociation System (Worthington Biochemical, LK003153). Dissociated cortical neurons were cultured in poly-D-lysine (Thermo Scientific, A3890401) and laminin (Sigma-Aldrich, L2020) coated slides and maintained in neurobasal plus (Thermo Scientific, A3582901) medium supplemented with B-27, 1% Glutamax, and 1% penicillin-streptomycin at 37 °C and 5% CO2. Neurons were treated with recombinant EPHRINB2 protein (R&D Systems, 496-EB-200) at a concentration of 1ug/ml at days in vitro (DIV) 2. Neurons were stained with MAP2 (Abcam, ab5392, 1:5000) at DIV5. Images were acquired using Leica DMi8 microscope and analyzed using Neurolucida 360 (MBF Bioscience).

### Cortical progenitor isolation

E15.5 embryo brains were extracted and cortices were dissociated into single-cell suspension using Neural Tissue Dissociation Kit (P) (Miltenyi Biotec, 130-092-628). Progenitors were magnetically selected and isolated using Anti-Prominin1 MicroBeads (Miltenyi Biotec, 130-092-333). Following progenitor isolation, total RNA was extracted and processed for RNA library preparation and sequencing.

### Postnatal neuron enrichment

Cortices were dissected from P1 and P14 brains and dissociated into single-cell suspension using Neural Tissue Dissociation Kit (P) (Miltenyi Biotec, 130-092-628 for P1 and 130-094-802 for P14). Cortical neurons were enriched using a neuron isolation kit (Miltenyi, 130-115-389). Following neuron enrichment, total RNA was extracted and processed for RNA library preparation and sequencing.

### RNA library preparation and sequencing

RNA sequencing was performed as previously described (103–105). Total RNA was extracted from *Zmiz1*^*fl/fl*^ and *Zmiz1*-KO samples using GeneJET RNA Purification Kit (Thermo Fisher, K0732). RNA concentration and RNA integrity (RIN) number were determined using Qubit RNA High Sensitivity Assay Kit (Thermo Fisher, Q32852) and Bioanalyzer RNA 6000 Nano assay kit (Agilent, 5067-1511). The RNA library was prepared using TruSeq RNA Library Prep Kit v2 (Illumina, RS-122-2001) according to the manufacturer’s instructions. The mRNA library was quantified using Qubit dsDNA High Sensitivity Assay Kit (Thermo Fisher, Q32851) and verified using Bioanalyzer DNA1000 assay kit (Agilent, 5067-1505). Verified libraries were sequenced with either NextSeq 500/550 High Output Kit v2.5 (150 Cycles) (Illumina, 20024907) or NextSeq1000/2000 P2 Reagents (200 Cycles) v3 (Illumina, 20046811) on a Nextseq500/550 or Nextseq1000/2000 system respectively. Sequenced reads were aligned to the mouse (mm10) reference genome with RNA-Seq alignment tool (STAR aligner). Aligned reads were used to quantify mRNA expression and determine differentially expressed genes using the RNA-Seq Differential Expression tool (Deseq2) (version 1.0.1). Alignment and differential expression analysis were performed using tools in the Illumina BaseSpace Sequence Hub. Overrepresentation analysis of differentially expressed genes was performed using WEB-based Gene SeT Analysis Toolkit (WebGestalt)(106). E15.5 progenitors sequencing data have been deposited in the Gene Expression Omnibus (GEO) database with accession no. GSE225333. P1 and P14 enriched neuron sequencing data have been deposited in the Gene Expression Omnibus (GEO) database with accession no. GSE241140.

To review GEO accession GSE225333:

https://nam11.safelinks.protection.outlook.com/?url=https%3A%2F%2Fwww.ncbi.nlm.nih.gov%2Fgeo%2Fquery%2Facc.cgi%3Facc%3DGSE225333&data=05%7C02%7Crkc%40tulane.edu%7C5dd0dc45aa47493c075708dca65ab084%7C9de9818325d94b139fc34de5489c1f3b%7C0%7C0%7C638568155652645734%7CUnknown%7CTWFpbGZsb3d8eyJWIjoiMC4wLjAwMDAiLCJQIjoiV2luMzIiLCJBTiI6Ik1haWwi.LCJXVCI6Mn0%3D%7C0%7C%7C%7C&sdata=2ERy33xa3cyfWvvEjnj48HdfQ6dxlR08x%2FQHcvJGaWk%3D&reserved=0

Enter token kpmpaumypfuhzwn into the box. To review GEO accession GSE241140:

https://nam11.safelinks.protection.outlook.com/?url=https%3A%2F%2Fwww.ncbi.nlm.nih.gov%2Fgeo%2Fquery%2Facc.cgi%3Facc%3DGSE241140&data=05%7C02%7Crkc%40tulane.edu%7C7a1004e0f1284d03bc8608dca65ad19b%7C9de9818325d94b139fc34de5489c1f3b%7C0%7C0%7C638568156201655213%7CUnknown%7CTWFpbGZsb3d8eyJWIjoiMC4wLjAwMDAiLCJQIjoiV2luMzIiLCJBTiI6Ik1haWwiLCJXVCI6Mn0%3D%7C0%7C%7C%7C&sdata=ned4ygoKVAOLPmIy6bnBksKD%2FlFlt31IuwDED8Mia3s%3D&reserved=0

Enter token klyhaosmvlmdrcx into the box.

### Chromatin sequencing immunoprecipitation (ChIP)

Cell culture: Neuro-2a (N2a) (ATCC, CCL-131) cells were cultured in DMEM-high glucose (Cytiva, SH30022) supplemented with 10 % FBS (Cytiva, SH3008803) and 1% Penicillin-Streptomycin (Thermo Fisher, 15070063) at 37 °C and 5% CO2.

*Zmiz1* cloning: *Zmiz1* cDNA was amplified using the following primers.

ZMIZ1_Forward: ATGAATTCTATGGACAGGCAC,

ZMIZ1_Reverse_HA: TCAAGCGTAATCTGGAACATCGTATGGGTAGTTGTTCTCAAAGAGAGACAG

EcoRV restriction sites were added to both primers and HA sequence was added to the reverse primer. pcDNA3.1 was digested using EcoRV and amplified *Zmiz1* cDNA was cloned into pcDNA3.1 vector. pcDNA3.1-Zmiz1-HA (*Zmiz1*-HA) clone was sequence verified and overexpression was confirmed by western blot.

ChIP: *Zmiz1*-HA plasmid was transfected into N2a cells and subsequently, ChIP DNA samples were obtained by using ChIP-IT Express Enzymatic Kit (ActiveMotif, 53009) following manufacturer instructions. Briefly, cells were fixed and crosslinked using formaldehyde and disuccinimidyl glutarate (DSG). Chromatin was enzymatically sheared followed by chromatin immunoprecipitation (IP). Anti-HA antibody (Cell Signaling, 3724S) was used for IP. IP DNA was quantified using Qubit DNA HS kit (Thermo Fisher, Q32851). The sequencing library was prepared using TruSeq ChIP Library Preparation Kit (Illumina, IP-202–1012). 10 ng of IP DNA per sample were used for library preparation. Library quality and quantity were assessed using Agilent DNA 1000 chip (Agilent 5067-1504) and Qubit DNA HS kit respectively. Libraries were sequenced using the NextSeq1000/2000 P2 Reagents (200 Cycles) v3 (Illumina, 20046812) on a Nextseq1000/2000 system. Sequencing analysis was done using the ChIPSeq App from BaseSpace Labs (Illumina), which uses MACS2 for region enrichment and HOMER for motif analysis. ChIP-Seq data were deposited in the National Center for Biotechnology Information’s Gene Expression Omnibus (GEO) database with accession no. GSE241255.

Go to https://nam11.safelinks.protection.outlook.com/?url=https%3A%2F%2Fwww.ncbi.nlm.nih.gov%2Fgeo%2Fquery%2Facc.cgi%3Facc%3DGSE241255&data=05%7C02%7Crkc%40tulane.edu%7C19fc84fba3904a07c80208dca65ae57a%7C9de9818325d94b139fc34de5489c1f3b%7C0%7C0%7C638568156521916142%7CUnknown%7CTWFpbGZsb3d8eyJWIjoiMC4wLjAwMDAiLCJQIjoiV2luMzIiLCJBTiI6Ik1haWwi.LCJXVCI6Mn0%3D%7C0%7C%7C%7C&sdata=Ys8NuDxQBIGqkAhrwcgeMsiXbFDIhltHG527gfn5%2BHw%3D&reserved=0

Enter token kxapqygsdpedlyx into the box

### Luciferase reporter assay

Sequences of the ChIP peaks were cloned into pGL4.20-Firefly luciferase plasmid (pGL4.20[luc2/puro]; Promega) using HindIII restriction site to generate pGL4.20-ChIPPeak-luc. Cloning was verified by DNA sequencing. pGL4.75 [hRluc/CMV] vector (Renilla reniformis) was used as expression control. Neuro-2a cells were transfected with a pool of control non-targeting siRNA (Horizon, D-001810-10-05) or *Zmiz1* siRNA (Horizon, L-007034-00-0005) using Lipofectamine3000 (Thermo Fisher, L3000015) and 24 hours later, cells were transfected with pGL4.20-ChIPPeak-luc and pGL4.75 (6 independent biological replicates). Luciferase activities were assayed using the Dual-Luciferase Reporter Assay System (Promega). Firefly luciferase expression was normalized with renilla luciferase expression for each condition and percentage luciferase activity was calculated relative to the baseline activity of control siRNA treated pGL4.20-ChIPPeak -luc. The final concentration of siRNA solution was 200 nM.

ChIP Peaks cloned:

#### Auts2

>mm10_dna range=chr5:133140936-133140986

GTGTGTGTGTGTGTGTGTGTGTGTGTGTGTGTGTACATGTGTATAGGTATG

>mm10_dna range=chr5:132601414-132601464

ACACATACATACACACATGCACACACACACATACA CACACATGCACACACA

>mm10_dna range=chr5:132568611-132568661

TGTGTTTGTCTCTCTCTCTCTCTTTCTCTCTCTCCC TCCCTCCCTCCTTTC

#### Epha5

>mm10_dna range=chr5:84383968-84384018

CACACACACACACTCACACACACACAGAGAGAGA GAGAGAGAGAGAGAGAG

>mm10_dna range=chr5:84347365-84347415

GCAGTCATCTTTGTTTCTATTTCTCTGTCTCACTCT TTCAGGGGAGTGTGT

>mm10_dna range=chr5:84085158-84085208

ACACACACACACACACACACACACACACACACACA CACACACAGAGAGAGA

#### Unc5d

>mm10_dna range=chr8:29497721-29497771

ACACAGAGACACACACAGACACATGTACACATACA GACACATGTACACATA

>mm10_dna range=chr8:29077995-29078045

GTGTGTGAGAGAGAGAGAGAGAGAGAGAGAGAG AGAAAGAGAGAGAGAAGG

>mm10_dna range=chr8:28721674-28721724

GGAGATAGAGAAAGAGACAGACAGAAACAGAGAG CCAAAGGTAGAGAGAGA

#### Efnb2

>mm10_dna range=chr8:7132492-7132542

TGTATATGTATATGTGTGTGTGTGTGTGTGTGTGT GTGTGTGTGTGTGTGT

>mm10_dna range=chr8:7885145-7885195

CTGTGTCTGGCAGTGTGTGTCTGTATATATCTCTC TGTGTGTGTCTCTGTG

>mm10_dna range=chr8:8167465-8167515 ACATGCACATAAATATACAAACATCCACACAGAAG CACATAGACATGCACA

#### Lhx2

>mm10_dna range=chr2:38341911-38341961

TGTGTGTGTGTGTTTCTTTCTCCCTCTCCCTC CTCTCTCTCTCTCTCTCTC

### Behavioral assay

Open Field test: All mice were of similar age (4-7 months) and weight. Locomotor exploration of an environment was conducted in an automated open field chamber of 16” x 16” (Accuscan Instruments, Columbus, OH) using methods previously described (107). Briefly, mice were allowed to acclimate to the testing room for 1 hour following transportation from the vivarium. Subjects were placed in the center of the chamber and recorded for 10 minutes. Automated built-in photobeams and overhead cameras recorded mouse behavior. Light intensity in the chamber was 100-200 lux and was isolated from external light. After testing, mice were placed in a new cage and did not interact with untested cage mates. The chamber was cleaned thoroughly with 70% ethanol and water between subjects. Locomotion and behaviors were recorded in the inner and outer zones of the chamber.

Elevated Plus Maze: All mice were of similar age (4-7 months) and weight. Anxiety evaluation was conducted in an elevated plus apparatus with 30 cm by 5 cm arms (closed arm height 15cm) elevated 65 cm above the ground as previously described in (108). Briefly, mice were acclimated for 30 minutes following transportation before testing. Light intensity in the maze was 250-350 lux, and the maze was isolated from external light or cues. Mice behaviors, including time in each arm, grooming bouts, head dips, and rears were recorded for 5 minutes with an overhead camera. BORIS event-logging software was used for analysis by an experimenter blind to genotype.

Ultrasonic Vocalizations: Performed as described in (109) on P4, P5, P7, P9, and P10 pups. Pups were isolated from their mothers for 5 minutes and placed into a glass beaker under a Pettersson M500-384 ultrasound microphone in a sound-proof chamber in a sound-proof room and recorded for five minutes. Recordings were analyzed in VocalMat (110).

Three Chamber Test: All mice were of similar age (4-7 months) and weight. Tests were performed as previously described in (111). The three-chamber apparatus was 60 cm by 43 cm by 22 cm, and each chamber was 20 cm wide isolated from external cues. Novel mice and subject mice were housed in separate rooms between tests, transported separately, and housed in different areas during testing to prevent any contact including olfactory. Mice acclimated for 1 hour before testing. Subject mice were first placed in the middle chamber with the doors to the side chambers closed and allowed to habituate for 10 minutes. Then the doors were opened, and the mouse was recorded for 10 minutes by an overhead camera. Behaviors were analyzed using BORIS event-logging software by an experimenter blind to genotype.

Prepulse Inhibition (PPI): All mice were of similar age (4-7 months) and weight. The PPI test was performed as described in (109) using an SR-Lab Startle response system (SD instruments). Mice were acclimated for 1 hour following transportation. To measure startle response, PPI, and habituation, we employed a three-block assay. Before the start of the tests, subjects were placed in the restraint tube, and white noise stimuli were presented. There was a 5-minute acclimation period in the chamber with a 70 dB background noise. The first block of the test consisted of six trials of a 120 dB startling stimulus presented for 40 ms. Block #2 was used to calculate PPI and consisted of twelve startle trials, three 76 dB prepulse trials, three 82 dB prepulse trials, three 85 dB prepulse trials, three 88 dB prepulse trials, and eight no stimulus trials. All trials contained a randomized 10–30 s gap between stimuli that averaged 20 s. No stimulus trials consisted of 100ms 70dB white noise only. Block #3 was identical to Block #1 (six, 120 dB alone startle stimuli) to assess for habituation and sensitization). Percent PPI was calculated as 100 minus 100 * (average startle amplitude with the prepulse/average startle amplitude of the Block#2 startle stimulus alone trials). Percent habituation/sensitization was calculated as 100 *((average startle amplitude of Block #1 minus average startle amplitude of Block #3)/average startle amplitude of Block #1).

Spontaneous Alternation T-Maze: All mice were of similar age (4-7 months) and weight. Test was performed as previously described in (107). The dimensions of the T-maze were 24 inches by 7 inches short arm with a closing gate, perpendicular to two 15 inches by 7 inches arms (left and right). Each mouse was transferred from their home cage to the starting position behind the closed gate. A complete experiment consisted of six trials. Latency to select the left or right arms was measured. If the mouse did not choose after three minutes in any trial, the experiment would end. Inter-trial interval of 30 seconds, after which the mouse was placed again in the start area. The accuracy in alteration was measured in five consecutive trials.

### Statistical analysis

Data analysis was performed using GraphPad Prism version 9.5.1 (www.graphpad.com) or using Matlab2023 (MathWorks). Data are presented as mean ± SEM. Two-tailed Student’s t-tests or ANOVA with Tukey correction for pairwise comparisons were used for statistical significance as indicated in the figure legends. P values < 0.05 are considered statistically significant. Asterisks indicate significance (*p<0.05, **p<0.01, ***p<0.001, ****p<0.0001). All experiments were performed with at least three biological replicates.

### Materials and correspondence

Correspondence and material requests should be addressed to Maria J. Galazo.

### Data Availability

Sequencing data are deposited in GEO and can be accessed at accession numbers GSE241140, GSE225333, and GSE241255.

