## Supplemental Figures for "Loss of *Zmiz1* in mice leads to impaired cortical development and autistic-like behaviors"

**Supplementary Figures**

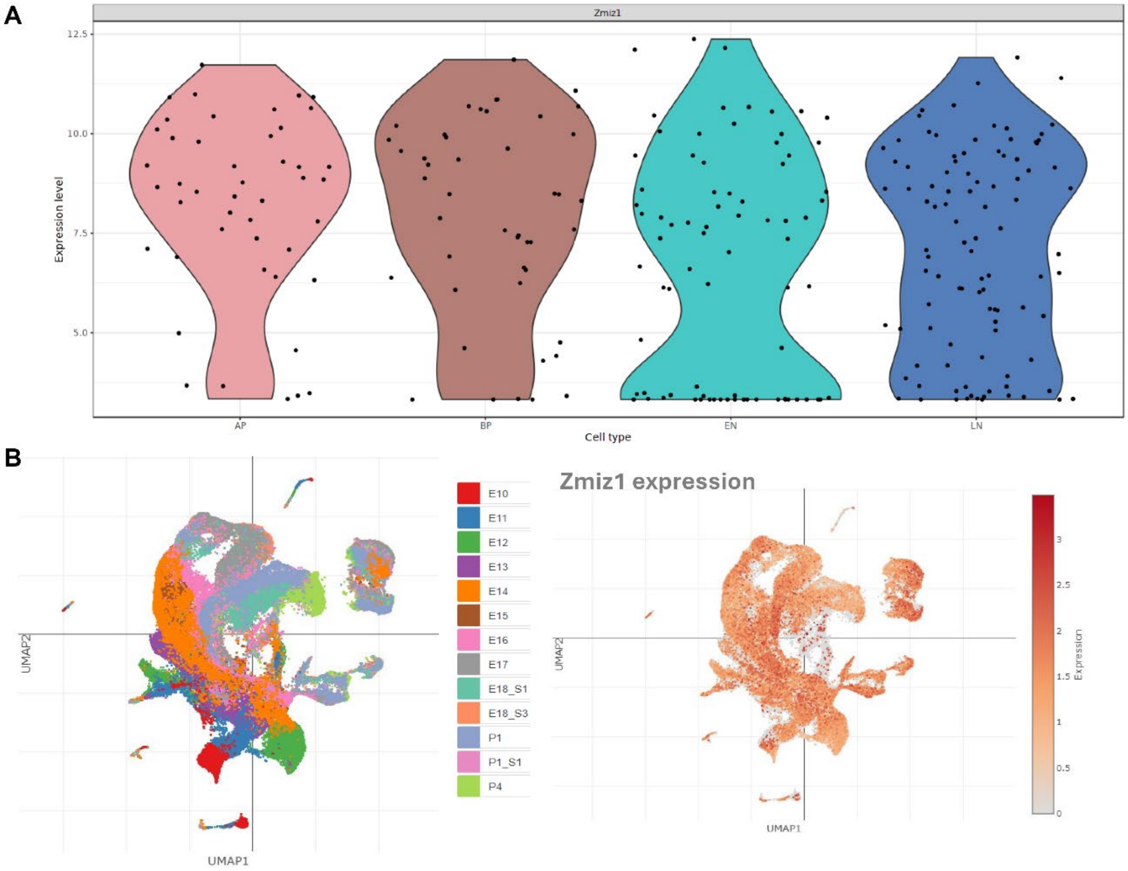


**Supplementary Figure 1: *Zmiz1* is expressed in neuronal lineages in the developing mouse brain**

(A) *Zmiz1* expression in the cortical progenitors and neurons. Adapted from http://genebrowser.unige.ch/science2016/ (16). (B) Temporal *Zmiz1* expression from E10 to P4 in the mammalian cortex. Adapted from Di Bella et al. 2021 (17). AP: Apical progenitors; BP: daughter basal progenitors; EN: Early neurons; LN: Late neurons.

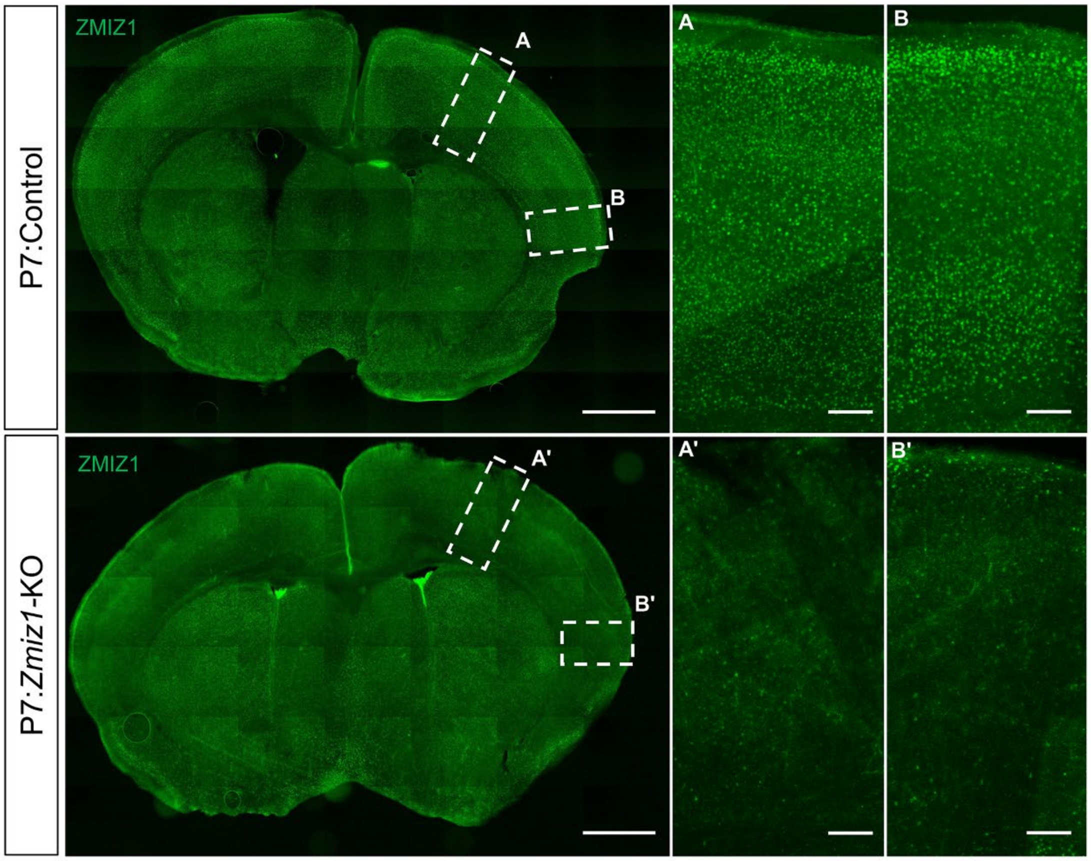


**Supplementary Figure 2: Efficient deletion of *Zmiz1* in *Emx1*Cre mouse model**

ZMIZ1 immunostaining in control and *Zmiz1*-KO brain depicting efficient ablation of *Zmiz1* in the *Zmiz1*-KO brain. Scale bar: 1 mm (left), 500 µm (A-B' insets). (A-A') Dorsal and (B-B') lateral cortical regions.

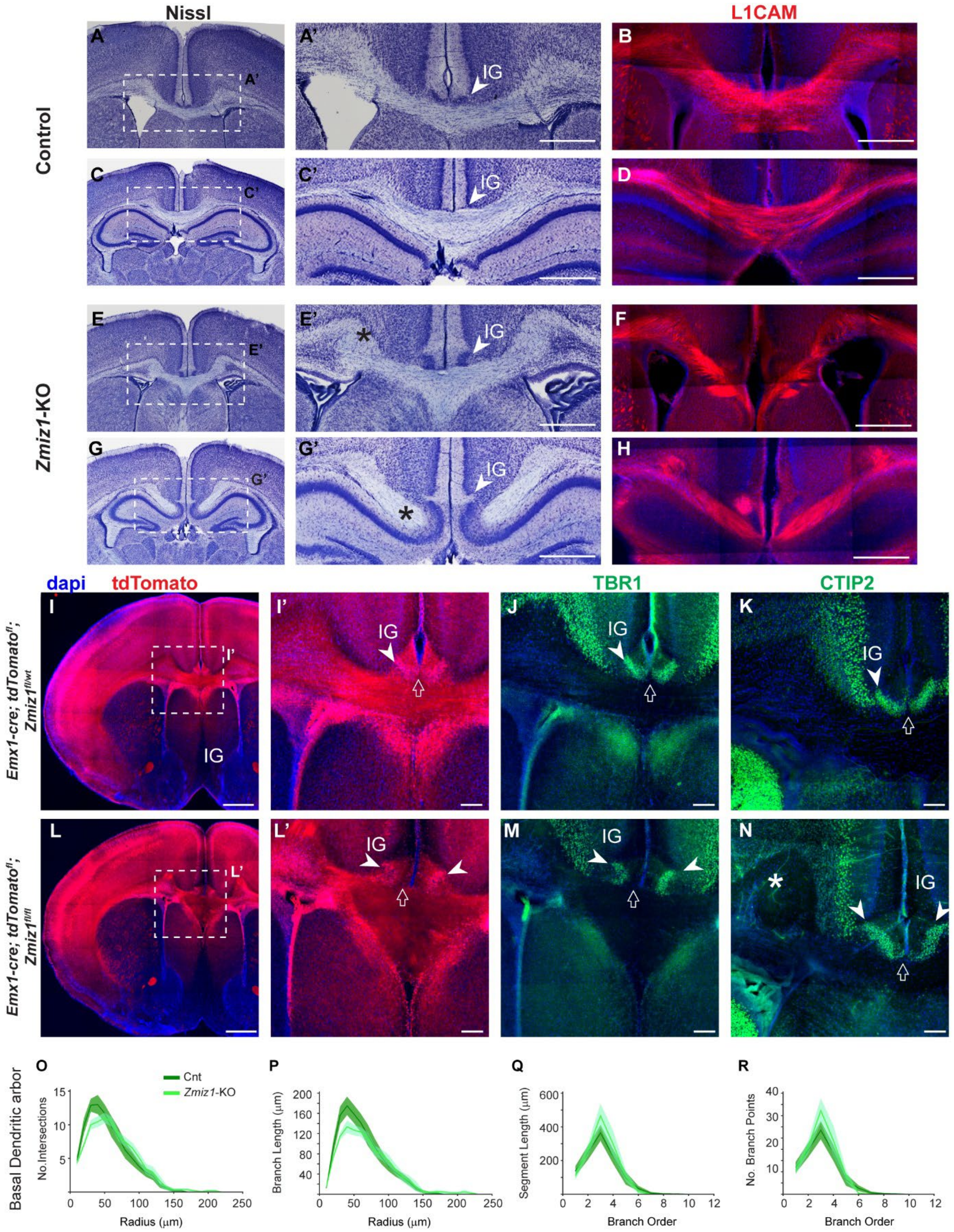


**Supplementary Figure 3: Formation of PB, alteration in IG positioning, and unaltered L2/3 basal dendritic arbors in *Zmiz1*-KO cortex**

(A, C, E, G) Nissl staining at P14 murine brain sections depicting alteration in Idusium griseum (IG) positioning in *Zmiz1*-KO mice. (A’, C’, E’, G’) Magnified views of dotted boxes in A, C, E, G. Asterisk indicates Probst bundle (PB), arrowheads show IG. (B, D, F, H) L1CAM staining reveals abnormal axon extension across the midline and corpus callosum in P14 *Zmiz1*-KO brains. (I, L) Abnormal midline structures and white matter were revealed using tdTomato reporter under control of the *Emx1*-Cre driver in P7 *Zmiz1*-KO brains. (I’, L’) Magnified views of dotted boxes in I, L. White-arrowhead points to IG, open arrow points to the dorsal cortical-CC boundary. (J, K, M, N) Defective boundary revealed by TBR1+ and CTIP2+ IG cell population in the *Zmiz1*-KO cortex. Asterisk indicates PB, arrowheads point to IG, arrows point to dorsal cortical-CC boundary. (O-R) Sholl analysis on L2/3 reconstructed neurons showing basal dendrites intersection by distance (O), basal dendrites length by distance (P), average branch length by branch order (Q), and number of branch points by branch order (R). Data are mean ± SEM. Two-tailed unpaired t-test showed no significant differences. Scale bar: 500 µm (A’, C’, E’, G’, B, D, F, H, I, L), 100 µm (I’, J, K, L’, M, N).

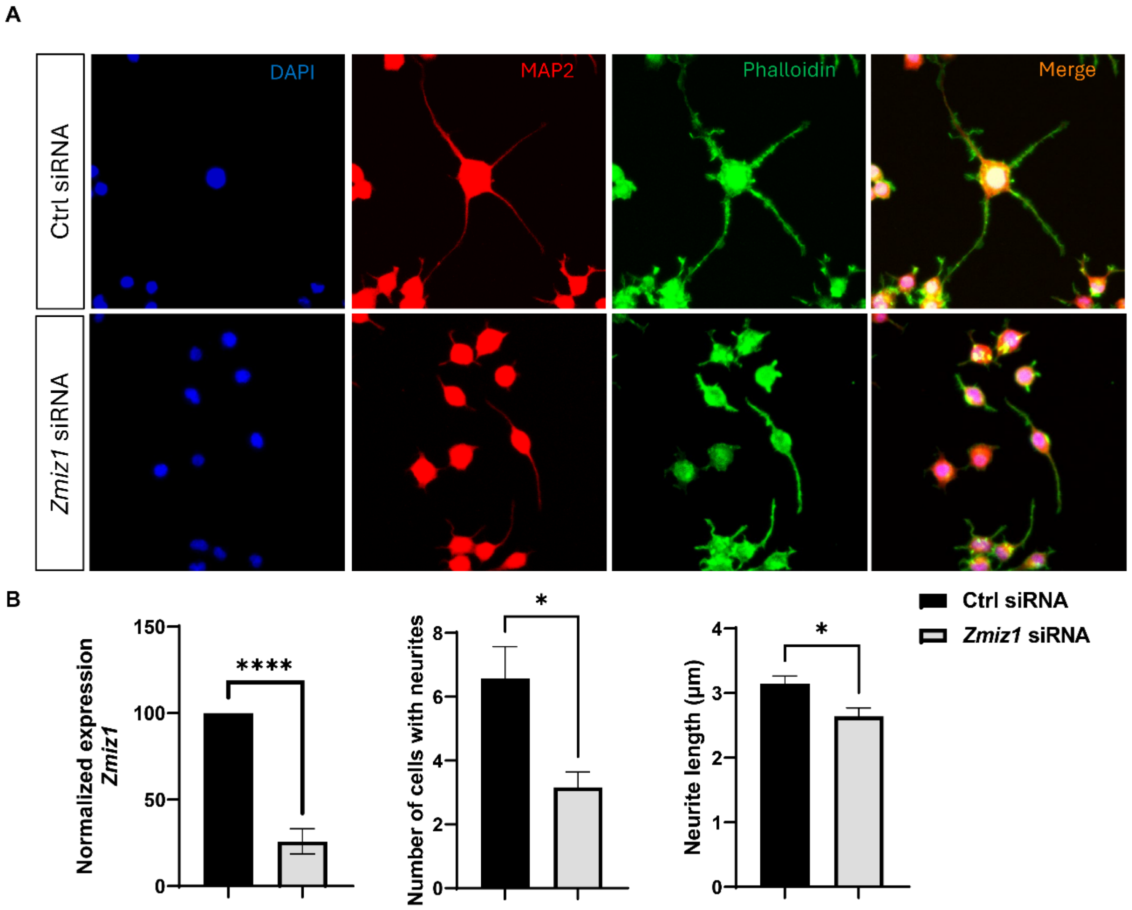


**Supplementary Figure 4: Deficit in neurite outgrowth in *Zmiz1* knockdown N2a cells**
(A) Visualization of neurites in differentiated N2a cells using MAP2 and Phalloidin stainings. Blue - DAPI, Red – MAP2, and Green – phalloidin. (B) Quantification of *Zmiz1* expression, neurite number, and length upon treatment with control and *Zmiz1* siRNA expression constructs. Scale bar: 50 µm. Data are mean ± SEM. * $P < 0.05$, two-tailed unpaired t-test. n = 3 biological replicates.

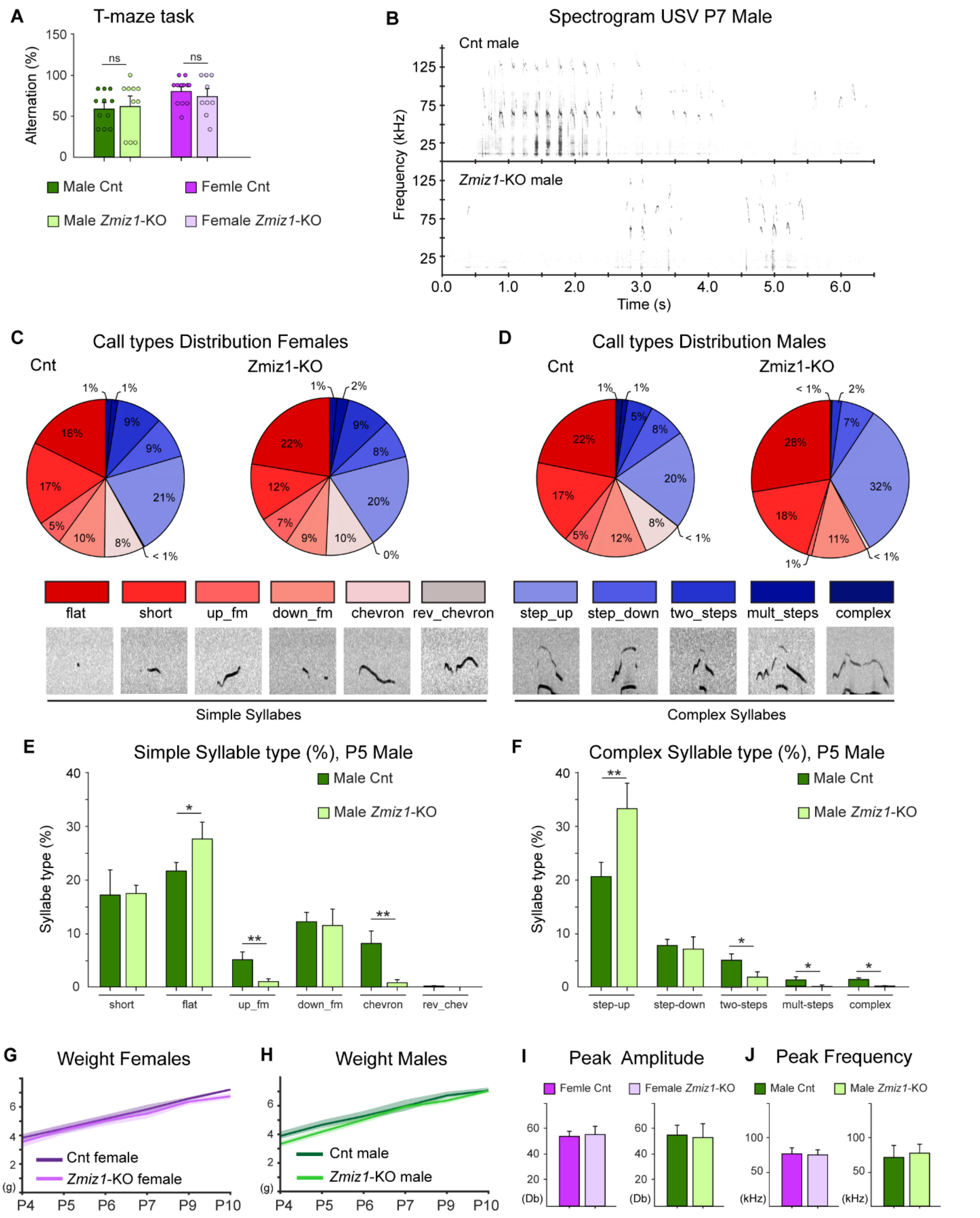


**Supplementary Figure 5: Altered USV in the neonates and no working memory deficits in *Zmiz1*-KO mice**

(A) T-maze behavioral test showed no difference in percent alternation. n = 12-15 per sex (control) and n = 12-15 per sex (*Zmiz1*-KO). (B) Representative USV spectrogram of P7 male. (C-D) Call type distribution across sex. *Zmiz1*-KO males produced less varied calls, but females exhibited no differences. (E) Simple syllable call types in P5 males. (F) Complex syllable call types in P5 males. (G-H) Charted weights of P4 to P10 female (G) and male (H) neonates used for ultrasonic vocalization (USV) test. (I-J) No difference in peak amplitude (I) and peak frequency (J) of USVs. Data are mean ± SEM. * $P < 0.05$, ** $P < 0.01$, ns – not significant, two-tailed unpaired t-test.

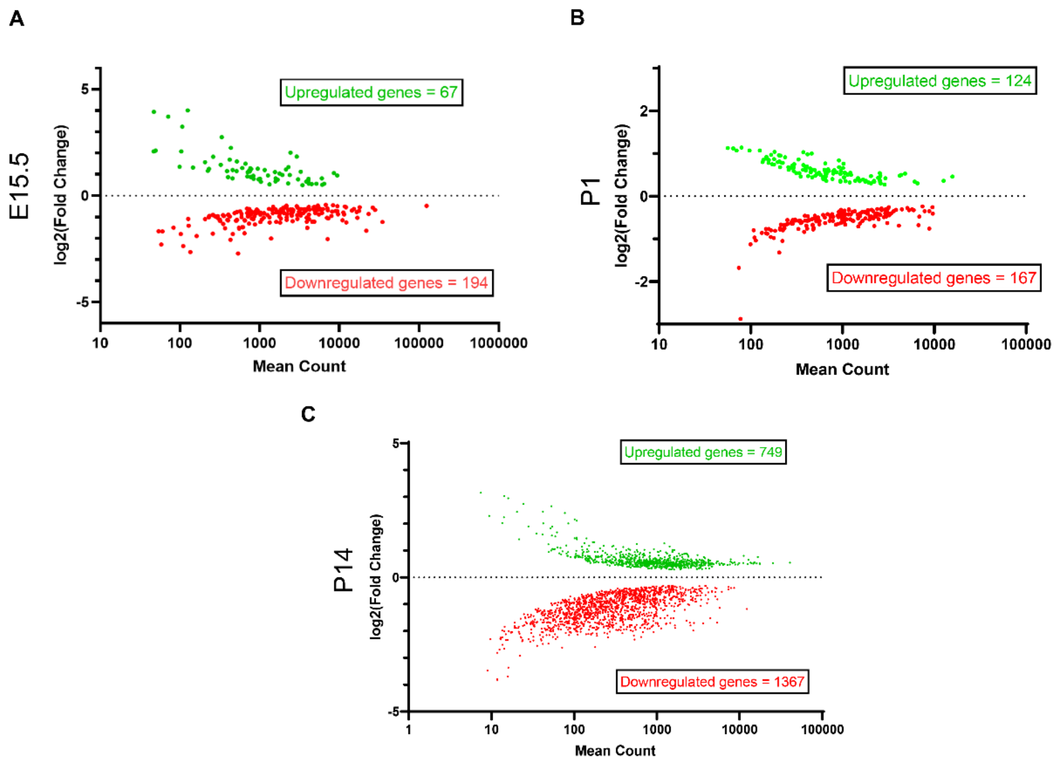


**Supplementary Figure 6: MA-plot of differentially expressed genes in the *Zmiz1*-KO cortex**

(A-C) MA-Plot of DEGs in *Zmiz1*-KO E15.5 progenitors (A), P1 enriched neurons (B), and P14 enriched neurons (C). Green – upregulated genes, red - downregulated genes.

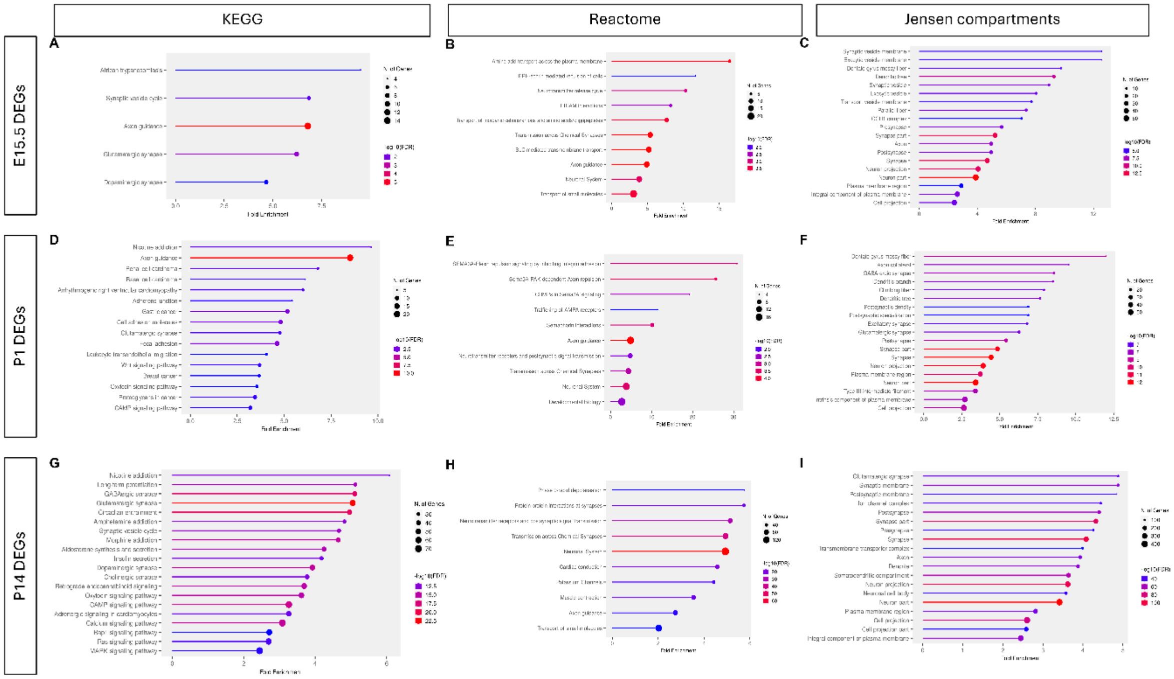


**Supplementary Figure 7: Gene ontology analysis of differentially expressed genes in *Zmiz1*-KO cortex compared to controls**

(A-C) Gene ontology analysis in E15.5 progenitors DEGs in *Zmiz1*-KO for KEGG (A), Reactome pathways (B), and Jensen compartments (C). (D-F) Gene ontology analysis in P1 enriched neurons DEGs in *Zmiz1*-KO for KEGG (D), Reactome pathways (E), and Jensen compartments (F). (G-I) Gene ontology analysis in P14 enriched neurons DEGs in *Zmiz1*-KO for KEGG (G), Reactome pathways (H), and Jensen compartments (I).

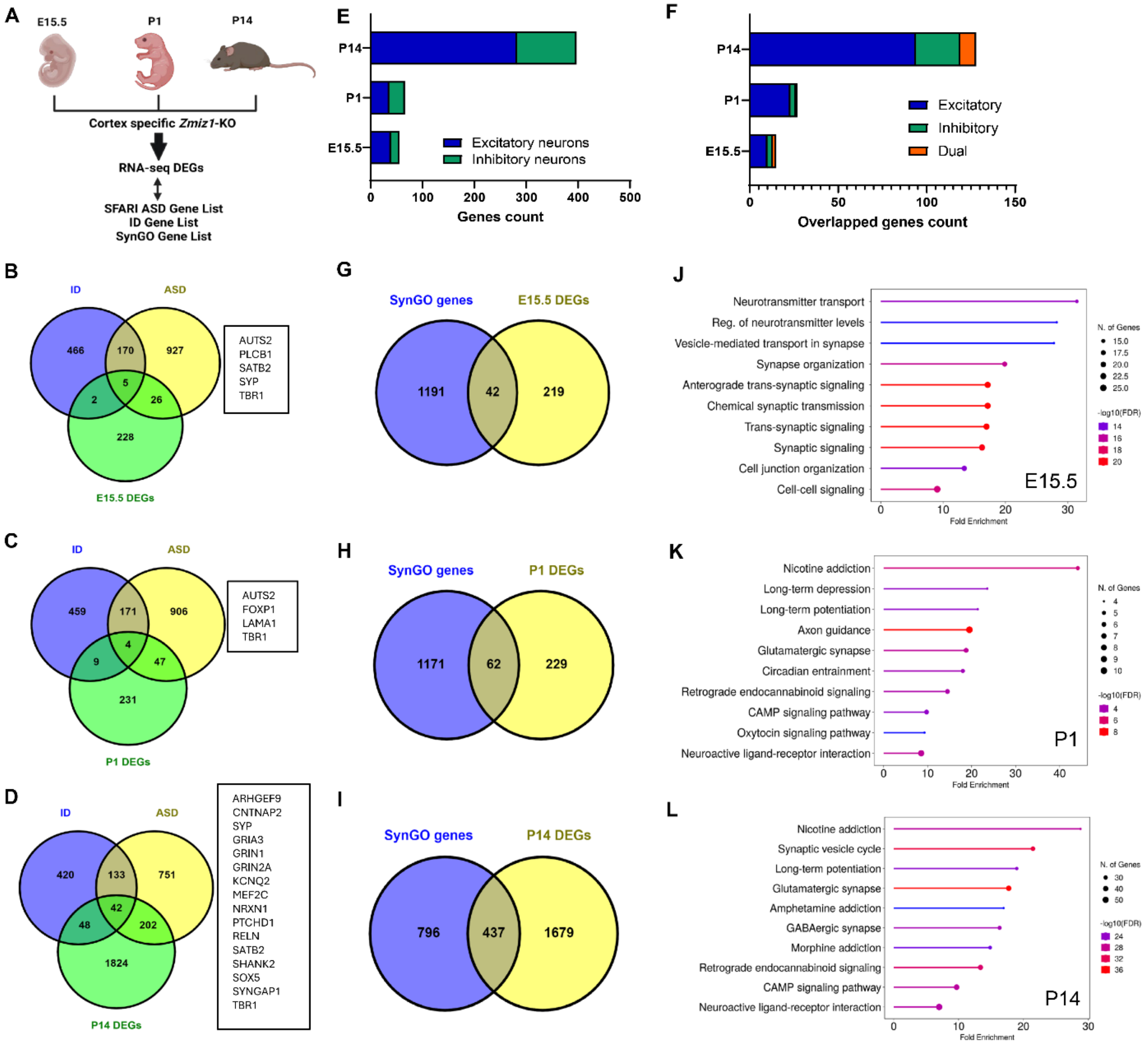


**Supplementary Figure 8: Abnormal regulation of synaptic-related and ASD/ID-risk genes in *Zmiz1*-KO cortex**

(A) Experimental approach showing E15.5, P1, and P14 DEGs mapping to SFARI, ID, and SynGO gene lists. (B-D) Venn diagrams showing overlap between E15.5 DEGs (B), P1 DEGs (C), and P14 DEGs (D) with ASD genes and ID genes. (E) Classification of E15.5 DEGs, P1 DEGs, and P14 DEGs into excitatory neurons and inhibitory neurons. (F) Classification of E15.5 DEGs, P1 DEGs, and P14 DEGs into excitatory, inhibitory, and dual synaptic genes. (G-I) Venn diagrams show overlap between E15.5 DEGs (G), P1 DEGs (H), and P14 DEGs (I) with SynGO genes. (J-L) ShinyoGO plot showing gene ontology for biological processes on E15.5 DEGs (J), P1 DEGs (K), and P14 DEGs (L) mapped to SynGO synaptic genes.

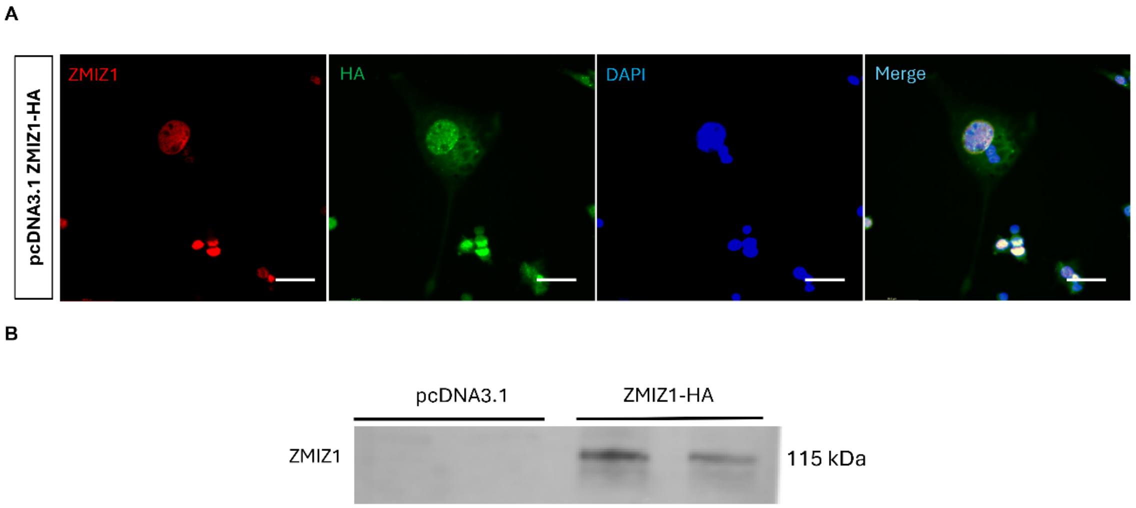


**Supplementary Figure 9: ZMIZ1-HA overexpression in N2a cells**

(A) Immunofluorescence antibody staining for ZMIZ1 (Red) and HA (Green) in N2a cells. Blue – DAPI, Scale bars; 10 µm. (B) Western blot for ZMIZ1-HA using the HA antibody.

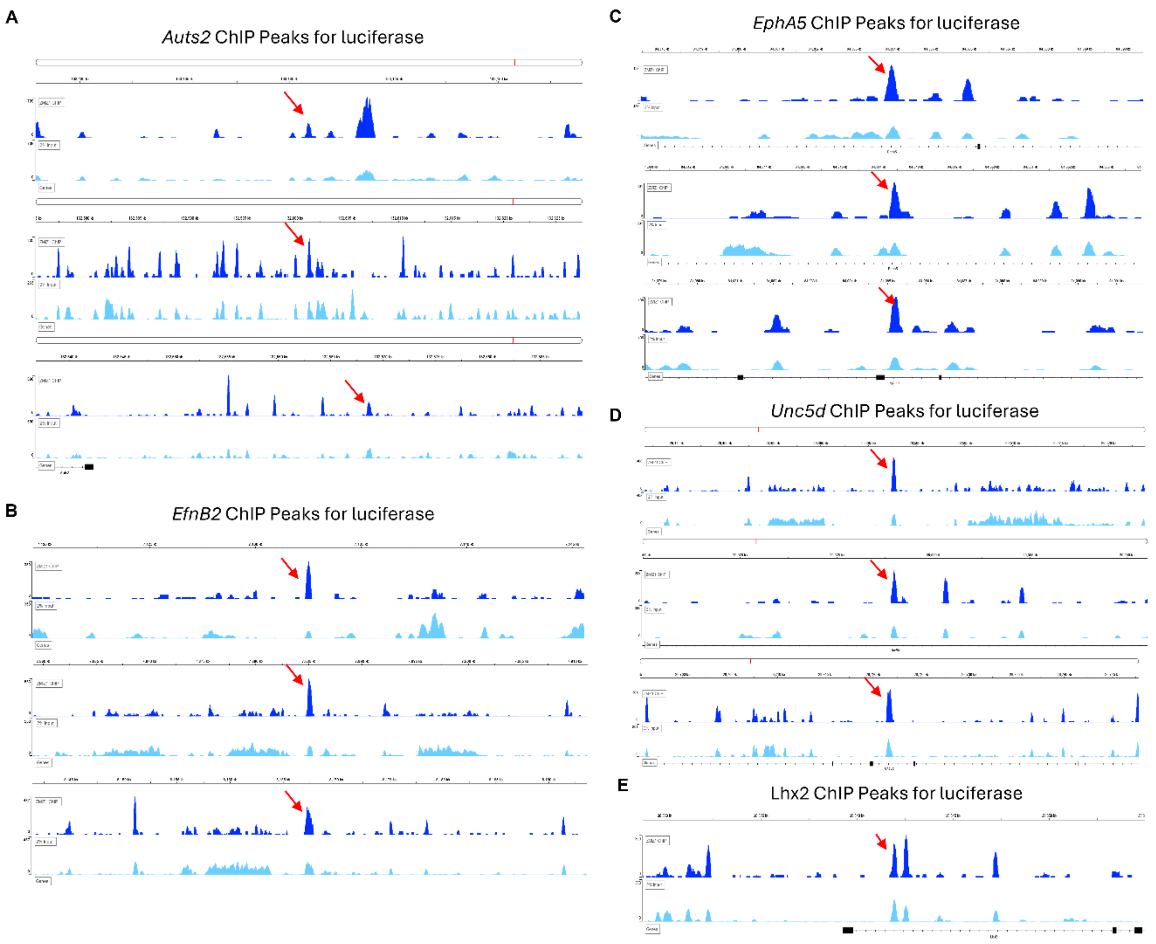


**Supplementary Figure 10: ZMIZ1 ChIP peaks used for cloning and luciferase assay**

(A-E) ZMIZ1 ChIP peaks for *Auts2* (A), *EfnB2* (B), *EphA5* (C), *Unc5d* (D) and *Lhx2* (E). Dark blue: ZMIZ1 ChIP, light blue: 2% input. Red arrows indicate ChIP peaks used for luciferase assays.

**A**

**Zmiz1 Motif**

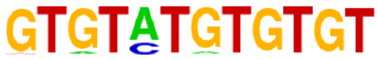


**B**

**Axon Guidance/neuron differentiation genes**

| | ChIP Seq. | | | RNA Seq. | |
|---|---|---|---|---|---|
| Gene Name | Annotation | No. of Peaks | Peak Score | E15.5 Fold Change | P1 fold change |
| Epha5 | Intron | 3 | 11.43349 | -0.68 | -0.49 |
| Efnb2 | Intergenic | 7 | 10.99742 | -0.7 | -0.3 |
| Lhx2 | Intron | 1 | 4.45808 | -0.77 | -0.52 |
| Auts2 | Intergenic | 4 | 12.33699 | -0.55 | -0.32 |
| Unc5d | Intron, Intergenic | 7 | 7.31162 | -0.84 | -0.49 |
| Slit2 | Intergenic | 1 | 4.59448 | 0.66 | -0.68 |

**Supplementary Figure 11: ZMIZ1 target motif and target regulation**

(A) ZMIZ1 top motif sequence from ChIPseq. (B) Overlapped genes in ChIP peaks, E15.5 DEGs and P1 DEGs showing gene name, number of ChIP peaks, and fold change at E15.5 and P1.
